# An age-depth model and revised stratigraphy of vertebrate-bearing units in Natural Trap Cave, Wyoming

**DOI:** 10.1101/2021.07.30.454390

**Authors:** David M. Lovelace, Cory M. Redman, Thomas A. Minckley, Blaine W. Schubert, Shannon Mahan, John R. Wood, Jenny L. McGuire, Juan Laden, Kathleen Bitterman, Holly Heiniger, Lindsey Fenderson, Alan Cooper, Kieren J. Mitchell, Julie A. Meachen

## Abstract

Almost a half-century ago excavations at Natural Trap Cave (NTC) began to yield evidence of the steppe paleoecology along the western slope of the Bighorn Mountains in north central Wyoming. The first decade of fieldwork led to the discovery of a diverse fauna that existed at the end of the Last Glacial Maximum. Stratigraphic deposits below the entrance of the cave were studied soon after excavations began, but never formally published. Although stratigraphy, taphonomy, and depositional circumstances were briefly discussed over the following years, little has been done to correlate the numerous stratigraphic schemes used by various authors. In this study, four stratigraphic sections were measured and analysed to establish an easily modifiable lithostratigraphic system of nomenclature. We provide the first correlations of all stratigraphic nomenclature used throughout excavations at NTC to facilitate comparisons with current and previous collections and publications. By leveraging more than 100 radioisotopic dates we developed an age-depth model and chronostratigraphic framework to further interrogate spatiotemporal relationships between strata, paleoenvironmental proxies, and fossil assemblages. Deposition is shown to be discontinuous; sediment accumulation in the study area is restricted to the buildup through peak penultimate and Last Glacial maxima. More recent (<10 ka) Holocene deposits unconformably cover the eroded surface of underlying Pleistocene strata. There is active reworking of sediments with transport and deposition of reactivated sediments within the Lower Chamber. We note that the two hiatuses coincide with interglacial periods and may reflect changing depositional circumstances within the cave such as extended periods of non-deposition, erosion, or bypass (possibly leading to deposition in the Lower Chamber). Contrary to previous reports, we demonstrate that it is unlikely a prominent snow cone existed or contributed to the pattern of sediment and fossil distribution within the study area, furthermore, we do not observe a continuous Pleistocene-Holocene transition in the study area. Further stratigraphic work will be needed to better understand the interrelationship between Main and Lower chamber deposits and the evolution of sediment accumulation in NTC.

## 1. Introduction

Upper Pleistocene sediments in the Rocky Mountain West record environmental and ecological histories during the last major advance and retreat of continental and alpine glacial ice (Illinoian–Wisconsin and Bull Lake–Pinedale glaciations; Nelson et al., 1979, Pierce, 2003; Benson et al., 2005; Pierce et al., 2018). Changing climate across this time period altered ecozones and shifted ranges for flora and fauna, leading to extirpation or extinction of the majority of vertebrate megafauna and establishment of non-analog floras and faunas (Graham and Grimm, 1990; Stafford et al., 1999; Koch and Barnosky, 2006; Stewart, 2009; Kohn and McKay, 2012). Studying these evolutionary events is often hindered by taphonomic biases (e.g., hydrodynamic sorting, variable deposition, time-averaging, and rates of accumulation) that make community level sampling more difficult (Olsen, 1980). High-resolution stratigraphic and temporal control can limit (but not eliminate) biases, although novel circumstances such as traps and ashfalls can minimize these biases and more faithfully capture signals of local ecology (Rogers and Kidwell, 2007). Many Late Pleistocene fossil localities in North American preserve diverse vertebrate assemblages (Kurtén and Anderson, 1980), and some sites are particularly well known for dense accumulations and/or diversity, for example: Rancho La Brea, CA; the Mammoth Site, SD; and the Ziegler Reservoir fossil site, CO (e.g., Agenbroad and Mead, 1994; Akersten et al., 1983; Sertich et al., 2014). These fossil deposits provide an extraordinary opportunity to answer far ranging questions about the evolution and extinction of local flora and fauna.

Natural Trap Cave (NTC) is another significant fossil locality, known for its exceptionally well preserved Late Pleistocene vertebrate fossils (e.g., Martin and Gilbert, 1978a & 1978b; Gilbert and Martin, 1984; Wang, 1988; Chorn, 1988). The cave contains fossils that span from late Illinoian to recent (i.e., ca. 150,000 ka - present); however, the vast majority of pre-Holocene specimens coincided with the last Pinedale glacial maximum (∼23,400 to 29,200 cal yr B.P.; Pierce, 2003), which terminated around ∼17,000 ka (Benson et al., 2005).

The fossiliferous deposits of NTC were first collected in 1969 (Rushin, 1973), subsequent excavations occurred throughout the 1970s and into the early 1980s (Martin and Gilbert, 1978a; Gilbert and Martin, 1984; Wang and Martin, 1993; Meachen et al., this issue); NTC was heavily sampled during this 11-year period (Wang and Martin, 1993; Meachen et al., this issue). Over the course of three years (1976-1978), geologist John Albanese developed seven cross sections, a description of sediments (including compositional analyses), and a working stratigraphic nomenclature for the cave (Albanese. 1976 Unpublished Report).

Although subsequent authors have drawn extensively from Albanese’s unpublished data (with some modifications in nomenclature including a reversal of the original numbering scheme; e.g., Martin and Gilbert, 1978a; Gilbert and Martin, 1984; Wang and Martin, 1993; Williams, 1993), no detailed stratigraphic work of the cave has been published to date. Furthermore, stratigraphic nomenclature was not consistently applied throughout the course of study at NTC, leading to confusion and difficulties in reconstructing previous excavations and placing collected material into a coherent stratigraphic and temporal context. Given the success of earlier excavations and the importance of a well-preserved repository of Late Pleistocene paleoenvironmental and paleoecological signals, a re-evaluation of stratigraphic and temporal relationships of deposits within NTC is warranted. After a 30-year hiatus, fieldwork at NTC resumed from 2014 to 2018 (Meachen et al., this issue), leading to new insights into paleobiogeography, depositional history, palynology, phylogeny, and an updated geochronology.

This paper aims to correct any confusion by establishing a correlation between early published and unpublished accounts of sedimentary deposits within NTC, as well as reconcile historic and recently acquired ^14^C determinations in order to construct an age-depth model. Our age-depth model provides a more comprehensive understanding of chronostratigraphic relationships and creates a more stable foundation upon which paleoecological inferences can be built, and previous hypotheses can be tested.

## 2. Geographic and historical context

Natural Trap Cave (NTC) is a karst pit cave located on the western slope of the northern Bighorn Mountains northeast of Lovell, Wyoming (Fig. 1) on a plateau (aka Little Mountain) at an elevation of 1,512 m (4960 ft). The cave entrance is situated on a narrow ridge that is sloping westward and flanked by deeply incised ephemeral surface drainages. It should be noted that the geographical depression containing the cave entrance represents a very limited catchment (< ca. 40,000 m^2^). In 1973, the Bureau of Land Management placed a steel gate across the 6 x 8.5 m horizontal entrance to protect cave resources and prevent accidental falls. A 3.5 m vertical drop below the gated entrance ends at a 2 m wide ledge that runs along the southwestern edge before transitioning to a final vertical drop into the main chamber where the current chamber floor is 24.5 m below the grate. The roughly bell-shaped main chamber is ∼43 m in diameter with a perimeter that is subcircular and uneven. The cave was originally mapped between 1969 and 1972 (Hill et al., 1976, p.86-87). In 2016 the cave was re-surveyed with improved techniques (see section 3.1 below).

**Figure 1.**
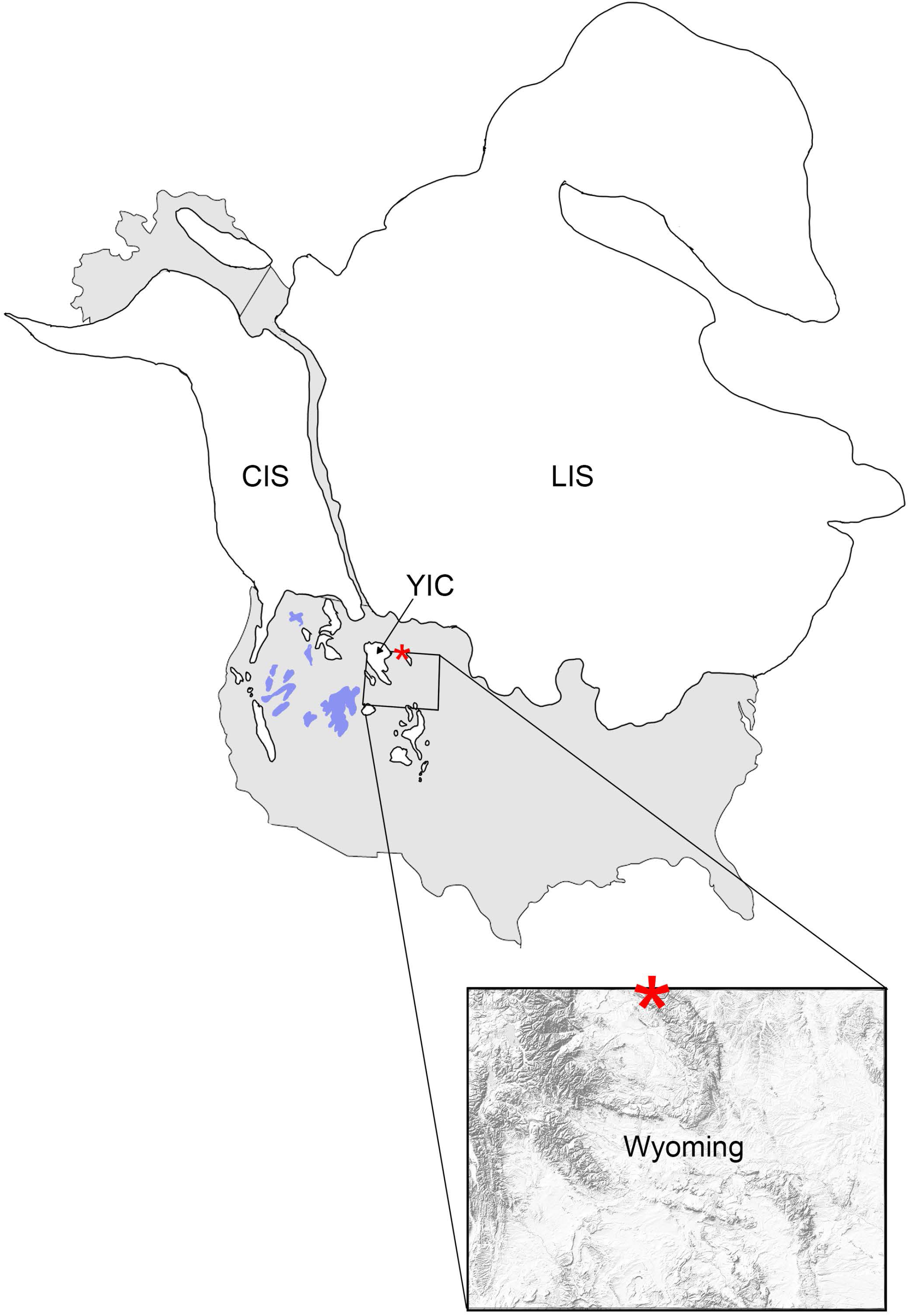
Geographic position of Cordilleran and Laurentide ice sheets (CIS, LIS, respectively), Yellowstone ice cap (YIC), Natural Trap Cave (asterix), and glacially influenced giant lakes (blue). Ice sheet reconstruction approximates ice boundaries as the Beringian corridor began to coalesce prior to ca. 23 ka ago (Heintzman et al., 2016; Batchelor et al., 2019)

The bedrock geology of Little Mountain plateau largely consists of the clastic-dominated Amsden Formation (Early Pennsylvanian; May et al., 2013) that overlies the Early-Middle Mississippian Madison Limestone Formation (Lageson et al., 1979; Sonnenfeld, 1996). The geologic setting of the cave, is an important component to the stories for the stratigraphy and the latest efforts of geochronology, and we refer the reader to Mahann et al., (this issue) for a more thorough examination of the importance of Amsden and Madison contributions to in-cave sediment.

Geographically, the cave is positioned south of a gap that developed between the Laurentide and Cordilleran ice sheets in central North America (Fig. 1) at the end of the Last Glacial Maximum (LGM) ∼15-14,500 cal ka BP (Dyke, 2004). This corridor was open prior to the coalescence of the two ice sheets >∼23 ka ago (Heintzman et al., 2016), and some taxa that spanned this passage would have been isolated north and south of the joined ice sheets for nearly 10,000 years. Subtle faunal interchanges between Beringia and North America are known to have occurred when this ‘ice-free corridor’ re-emerged uniting taxa divided by the coalesced ice sheets; NTC is ideally situated to capture that signal (Shapiro et al., 2004; Meachen et al., 2016; Pedersen et al., 2016; Heintzman et al., 2017; Wang et. al, 2021). Although the entrance to the cave was certainly known to the earliest people in the region (Crow Elder Grant Bulltail, Pers. Comm to JAM), only one known artifact has been found: a red-jasper biface that was surface collected immediately below the cave entrance. A putative atlatl shaft was reportedly found in a packrat midden near the surface (see Martin and Gilbert, 1978a), but this artifact was later determined to more likely be a coup stick and a relatively recent addition to NTC ( Meachen et al., this issue).

### 2.1 Land acknowledgment

As researchers studying the geologic history of NTC, we must acknowledge that we are visiting the ancestral lands of the Apsaalooké (Crow), Očhéthi Šakówiŋ (Great Sioux Nation), and Tséstho’e (Cheyenne), lands that were forcibly taken from them by white colonizers during a period of extensive genocide. NTC is situated between Medicine Mountain, and the Pryor Mountains, a region that remains culturally significant to many Indigenous people. We recognize the sovereignty of Indigenous people throughout the region whose history stretches millennia into the past and who are an active part of the present. Although we may employ western science to frame stories hidden in rock, we must not overlook Native ways of knowing the history of the region as well. There is much to share and much to learn.

## 3. Methods

Original fieldwork was used in conjunction with unpublished data such as field notes, collection data, field and analytical reports (i.e., Albanese. 1976 Unpublished Report), and other communications. These data were obtained from a variety of sources including the University of Kansas Biodiversity Institute Division of Vertebrate Paleontology, Bureau of Land Management, and personal collections. These data can be found at http://dx.doi.org/10.17632/f8frrskzpz.1, an open-source online data repository hosted at Mendeley Data ([dataset] Lovelace et al., 2021).

### 3.1 Cave surveys

The Vedauwoo Student Grotto initially surveyed NTC between 1968 and 1972 using the steel tape and Brunton compass method. Their finished cave map was published in a compilation of Wyoming caves (Hill et. al., 1976, p. 86-87). Members of the 2016 field party were granted permission by the BLM to re-survey the cave using more accurate mapping methods, including DistoX and LiDAR technology. The DistoX survey used a modified Leica laser distance meter to measure distance, azimuth, and vertical angle. The instrument was calibrated for local declination prior to the survey. The LiDAR survey used a Zeb REVO to scan the cave; scanning and post processing were conducted by H2H Geoscience Engineering, Troy, New York. The principal goal was to determine if alternate entrances into the main chamber existed, produce a volumetric reconstruction of the cave, and ensure all observed passages were mapped in more detail.

### 3.2 Stratigraphy

Two unpublished stratigraphic schemes were devised (and used extensively during early excavations) by S.A. Chomko/K.A. Lippencott and J. Albanese, an alphanumeric ‘zone’ and ‘unit/sub-unit’ based nomenclature, respectively (Appendix Figure 1). Martin and Gilbert (1978a) were the first to publish a simplistic profile outlining seven discrete stratigraphic levels. Their descriptions of each ‘stratum’ are minimal, and no correlation to previous unit nomenclature(s) used during excavation was provided. Gilbert and Martin (1984) published another schematic, but no unit designations or descriptions were included. Wang and Martin (1993) proposed several hypotheses related to the origin and taphonomy of fossiliferous cave deposits, but the stratigraphic nomenclature used was that of Albanese (1976 Unpublished Report). The complexity of the deposits below the entrance of NTC required us to correlate new observations with previous stratigraphic work to avoid confusion between studies; something that had not been attempted before this paper.

During the 2016 field season four stratigraphic sections were measured with some of the units traceable between each section. Fresh sediment was exposed in vertical profiles at each section in order to observe a non-weathered surface. Grain-size, texture, color, thickness, sedimentary features and lateral relationships were recorded. Photography was used to record the lateral continuity of units across several meters and along the transition between north-south and east-west trending walls. Work was conducted under Bureau of Land Management permit PA-13-WY-207.

### 3.3 Geochronology

#### 3.3.1 Tephrochronology

At least five discrete volcanic ash accumulations have been observed within NTC deposits over the course of study (1972-2017); however, the lateral and temporal relationship of some ash deposits remain poorly constrained. Most ashes appear as primary deposition and can be stratigraphically traced through the deposits, however there is evidence of some remobilization of ash within the cave. Albanese (1976 Unpublished Report) was the first to note the presence of an ash (unmapped) along the southwall of the 1976 NTC excavation and remarked that little had been done with the ash up to that point; that still remains the case and should be a focus for future work within the cave. As excavations continued between 1977 and 1983, three new ash deposits described as a white volcanic ash layer overlying two grey volcanic ash layers were reported (Gilbert [unpublished NSF report] see [dataset] Lovelace et al., 2021; Gilbert et al., 1980; Gilbert and Martin, 1984) from units discovered below the lowermost deposits of Martin and Gilbert (1978a). In an abstract, Gilbert and others (1980) provide a ca. 110 ka BP fission-track date for the stratigraphically youngest of those three ashes (Mahan et al., this issue). Vertebrate remains, including *Ovis*, *Equus*, *Lepus*, along with microvertebrate material (Gilbert, 1980 BLM Report [unpublished] see [dataset] Lovelace et al., 2021; Gilbert and Martin, 1984; Wang and Martin, 1993) were found below the undated lowermost ash.

During the course of recent fieldwork two ashes were encountered low in section, rather than three as noted by Gilbert and Martin (1984). The ashes were grey in appearance, and we make the assumption, in the absence of direct correlation with previous observations, that these represent the lower two grey ashes of Gilbert and Martin (1984). Two additional ashes were observed, one on the southwall at the surface of the cave deposits that is likely the ash referenced by Albanese (1976 Unpublished Report), and a 20 cm thick, but laterally limited (<1 m) ash accumulation on the north wall just below the Upper Red layer (described below).

The middle of the ‘lower three ashes’ has a reported ^40^Ar/^39^Ar date of 129.6 ±17.4 ka (Mahan et al., this issue), sanadine was absent from the lowermost ash sample but zircons were present in a grab sample taken for palynology. A ^238^U/^206^Pb age of 140 ± 7 ka was determined for the lowermost ash (Clementz et al., this issue) indicating fossil accumulations at NTC extend into the Bull Lake Glaciation (Pierce, 2003). These two dates are used to help temporally constrain the lowermost sediments observed in the excavation.

#### 3.3.2 Radiocarbon

Clear stratification is present in the cave, and several successive units contain datable vertebrate fossils (Martin and Gilbert, 1978a; Gilbert and Martin, 1984). This has provided the foundation upon which larger scale interpretations of timing of floral and faunal changes have been made (e.g., Gilbert and Martin, 1984; Wang, 1988; Wang and Martin, 1993). Specimens used for radiocarbon analyses were from one of two institutions: **UW**: University of Wyoming Geological Museum; **KUVP**: University of Kansas Biodiversity Institute Division of Vertebrate Paleontology.

**DICARB (and 2 unknown) Determinations 1973-1980:** Eight ^14^C dates were determined from NTC specimens between 1973 and 1980. The first ^14^C determinations from bone found within NTC were analysed in 1973, and later published as 10,920 ±300 yr BP and 12,770 ±900 yr BP (Martin et al., 1977). No information other than the ages were published and no original data pertaining to the analytical lab, specimens, methods or results have been found; these dates were later perpetuated (Martin and Gilbert, 1978a) or cited as ‘*Loendorf, personal communication*’ (Gilbert and Martin, 1984; Williams, 2009). Considering the lack of supporting information, analytical lab, or methods, we consider these two ages suspect. Martin and Gilbert (1978a) report a ‘17,620 yr BP’ and ‘20,170 yr BP’ age, also with no error or associated data. The ‘20,170’ age remains unverified, however, the ‘17,620’ age is likely analytical number DICARB 690, published with an associated error ±1490-1820 yr BP (Martin et al., 1979) but is later misidentified as DICARB 1690 (Gilbert and Martin, 1984). Another typographical error occurred again when the age determination 14,670 ±670/730 yr BP (DICARB 689; Martin et al., 1979) was misidentified as DICARB 1689 (Gilbert and Martin, 1984). The datasheets for DICARB 689 and 690 have not been found. Three additional radiocarbon ages were reported in Gilbert and Martin (1984): 17,870 ±230 yr BP (DICARB 1686), 20,250 ±275 yr BP (DICARB 1687), and 21,370 ±830/920 yr BP, the correct determination for DICARB 1689; [dataset] Lovelace et al. (2021).

Of the eight historical radiocarbon dates listed above only the last three dates have preserved datasheets with specimen information, however both DICARB 1689, and 1687 contain more than one specimen combined “at the sender’s request” to produce enough datable material ([dataset] Lovelace et al., 2021). We do not consider the mixing of two specimens a reliable method to obtain a radiocarbon date and we consider the ages of these two ‘mixed sample’ dates to be suspect. As such, DICARB 1687 (20,250 ±275 yr BP) is the only radiocarbon age from NTC that has been verified (e.g., a known datasheet exists) and is based on a single specimen with stratigraphic control prior to 2001. DICARB age determinations have been questioned (Reuther and Gerlach, 2005), though their incorporation into larger datasets used in Bayesian models may still have utility (Hamilton and Krus, 2018).

**OxA Determinations 2001-2005:** Forty-three AMS ^14^C ages were determined from NTC specimens at the Oxford Radiocarbon Accelerator Unit (ORAU) between 2001 and 2019. Twenty-two of those ages have been referred to in publications (Shapiro, 2004; McKay, 2008; Williams, 2009; Salis et al., 2020; Perri et al., 2021; Salis et al., 2021; Table S1). **LLNL Determinations 2015-2018:** Fifty-eight (including two inter- and two intralab replicate samples) AMS ^14^C ages were determined from NTC specimens at the Lawrence Livermore National Laboratory CAMS facility (LLNL; Table S1). **KCCAMS Determinations 2015-2018:** Four AMS ^14^C ages were determined from NTC specimens at the Keck Carbon Cycle AMS Facility University of California-Irvine (KCCAMS; Table S1). See Appendix for methods used for OxA, LLNL, and KCCAMS determinations.

### 3.4 Age-Depth Model

Stratigraphic units within the cave represent a complex history varying between periods of relatively continuous deposition, subtle to significant erosion, or relatively stable surfaces with low sediment accumulation. The contacts between stratigraphic units range from gradational to unconformable. Although clear boundaries exist and can be traced across many meters of excavated surfaces, the units themselves are non-uniform, varying in thickness with lateral changes in lithology and contact type. For this reason, we employ an age-depth model using R (R Core Team, 2017) with Bacon (Blaauw and Christen, 2011; R package ‘rbacon’ v. 2.5.3) based on the IntCal20 calibration curve (Reimer et al., 2020) and a simple procedure for correcting age-depth relationships between sections with high variability in unit parameters such as depth and thickness (Pelton et al., 2017).

Dated materials have been collected from NTC since the earliest excavations, but not all have had their stratigraphic position recorded in a way that allows for simple age-depth relationship determinations. The first ^14^C dates and tephrochronology from NTC (∼1977-1980) gave some indication that the cave’s depositional history extends at least to 110 cal ka BP (early Sangamonian) and it was proposed that there was relatively continuous deposition to the present (Martin and Gilbert, 1978a; Gilbert and Martin, 1984). This has provided the foundation upon which larger scale interpretations of timing of floral and faunal changes have been made (e.g., Gilbert and Martin, 1984; Wang, 1988; Wang and Martin, 1993). Of those early ^14^C determinations, only one of the eight dates has verified analytical results and is based on a single specimen. To test the hypothesis that NTC preserves a continuous record of deposition that spans from (at least) 110 cal ka BP to present, we leverage a series of ^14^C determinations (n=105; including 4 replicate and 79 previously unpublished determinations; Table S1) measured over the last two decades, as well as recent ^40^Ar/^39^Ar and ^238^U/^206^Pb dates for material older than 45,000 years to constrain boundaries within an age-depth model. Most of the dated material does not have strict stratigraphic control, but all dates can be constrained to one of 11 lithostratigraphic units defined below.

The model presented below (section 4.3) is based on 22 radiometric dates with strict stratigraphic control; 21 dates are from the same analytical lab. Two significant hiatuses are interpreted from the strata and are included in the model, the youngest hiatus defines the boundary between the Upper Beds and Primary Beds; it is a clear erosional surface that incises through the Primary Beds along the South Wall. The second hiatus was placed at an abrupt change in lithology that we infer to be the boundary between the relatively continuous deposition of the fine-grained dominated Lower Pebbly Grey bed and the overlying boulder-sized breakdown of the Maroon Blocks (269 cm depth, Reference Section). Through an iterative set of modelling runs, we determined that the age-depth models should not assume conformal sedimentation. Despite the depth of dated material collected from the cave, we screened dated materials for positional accuracy and provenience, allowing for extrapolation to younger and older unconformable boundaries (Blaauw and Christen, 2011). Based on our selection criteria, we chose 22 of the >100 radiometric dates as appropriate for age-model determinations. Those 22 samples have the strongest stratigraphic control, the remainder are limited to unit-level (e.g., their position *within* the unit is not well constrained). In order to make sure we were not significantly altering model results by excluding 81 dates (a suggestion outlined by Hamilton and Krus, 2018), we repeated the study with the inclusion of all NTC ^14^C determinations from the last two decades, regardless of lab, as well as DICARB 1687. Given the lack of stratigraphic control beyond unit-level for the majority of specimens, ^14^C determinations without strict stratigraphic control were randomly distributed within their respective units (rather than stacking the dates chronologically within each unit, which would reach beyond the information at hand). We independently modeled two discrete sections (i.e., Upper Beds through Primary and Rainbow Beds [with a hiatus at 40 cm], and Lower Beds [all strata below 269 depth in Reference Section]). Age-depth relationships were estimated using 7,000-8,000 iteration model runs using R (R Core Team, 2017) with Bacon (Blaauw and Christen, 2011; R package ‘rbacon’ v. 2.5.3) based on the IntCal20 calibration curve (Reimer et al., 2020). Our assumptions were that sedimentation is relatively constant over time, and that contacts between the stratigraphic units were of similar age regardless of horizontal position on the cave floor.

## 4. Results

### 4.1 Survey

The results of the DistoX survey conducted in 2016 demonstrated a strong similarity to earlier mapping done by the Vedauwoo Student Grotto (Hill et al, 1976, p. 86-87), though the recent survey produced a more detailed map with a number of cross sections and map profile views (Fig. 2; for complete map see Fig. S2). A LiDAR survey was also conducted to provide a volumetric model of the cave system ([dataset] Lovelace et al., 2021). Although several domes in the ceiling of the main chamber were explored, no additional entrances to the cave were discovered; however, further exploration is warranted to more fully understand the evolution of NTC.

**Figure 2:**
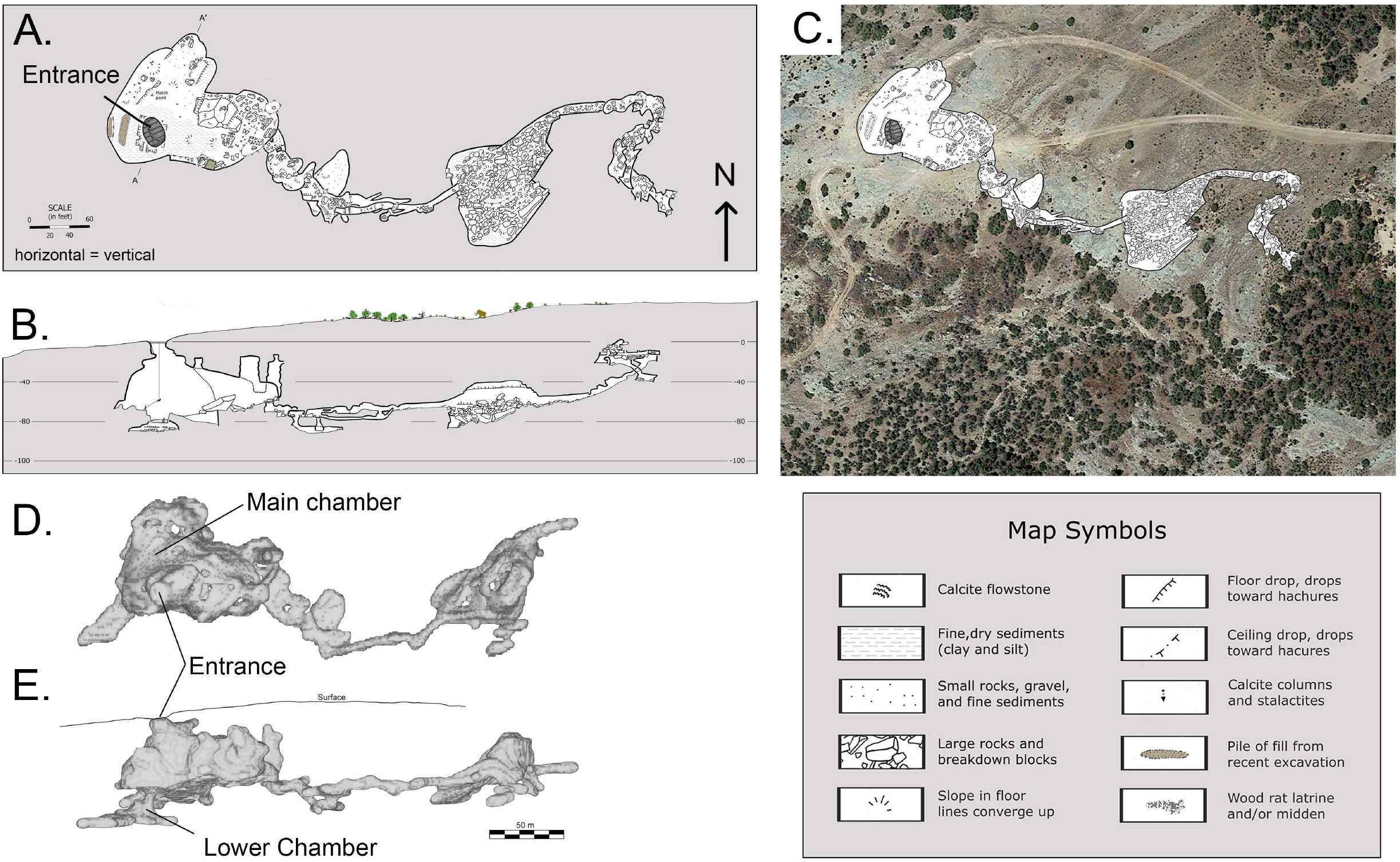
Plan (A;D) and profile (B;E) view of cave passages from DistoX (A-B) and LiDAR (D-E) surveys, with an aerial view (Google Earth) containing a superimposed cave map (C). Note: A-B scale in feet; modified after Hans Bodenhamer et al., 2016 (Fig. S2). D-E scale in meters ([dataset] Lovelace et al., 2021). Profile view (B) looking N20W.

During the 1985 excavation, an entrance into what is now called the Lower Chamber was breached in the deepest part of the main excavation (Meachen et al, this issue). Wang and Martin (1993) stated that the Lower Chamber contains Pleistocene fossils and it undoubtedly received sediment inputs from the overlying room where excavations have been focused. This was corroborated by brief forays into the Lower Chamber by members of the 2014 and 2016 field crews. Fifty-two vertebrate fossil specimens were collected from the surface of the Lower Chamber (2014: n=50; 2016: n=2; Appendix Table 1). An un-altered impression of a large bone was also observed. It is possible that this is the mammoth femur that was removed in 1985 along with a complete equid skull (Wang and Martin, 1993). Although the DistoX survey team did not survey this chamber, only part of the Lower Chamber was included in the LiDAR survey: further work is warranted to complete the survey of this portion of the cave system. The LiDAR survey of the Lower Chamber provides the only quantifiable information on the length, volume, and overall shape of this room and its spatial relationship with the overlying main chamber (Fig. 3).

**Figure 3.**
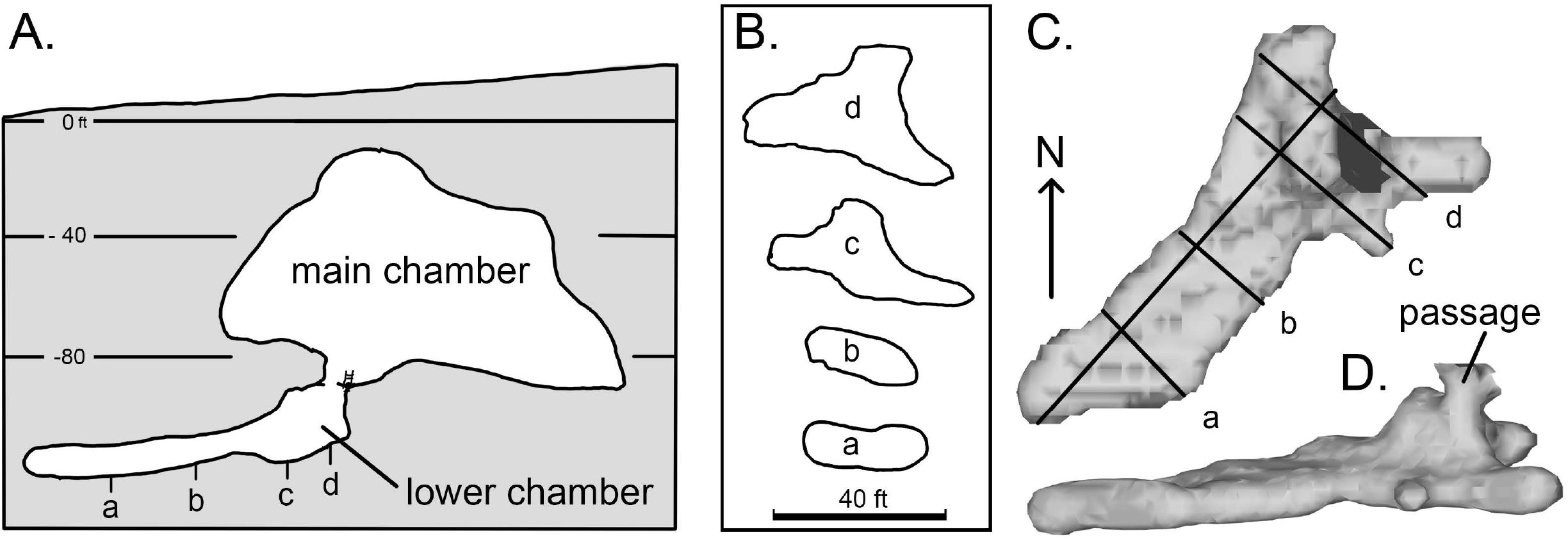
Cross section of main and Lower Chamber (A) ca. 5 m northwest of and parallel to A - A’ of Fig.MAP. Four cross sections of the Lower Chamber (a-d) show the constriction of the chamber downslope (B). 3D rendering of Lower chamber in map (C) and profile (D) view. Depth in feet.

**Table 1.** Stratigraphic description

| Specimen | Center | Center ID | Date | Error | to: | from: | % | Stratigraphic Interval | Depth |
| --- | --- | --- | --- | --- | --- | --- | --- | --- | --- |
| UW 53755 | LLNL | CAMS 176896 | 2,550 | 60 | 2375 | 2764 | 95.45 | Upper Beds | 15 |
| UW 51659 | LLNL | CAMS 179250 | 5,795 | 30 | 6499 | 6666 | 95.45 | Upper Beds | 25 |
| UW 53754 | LLNL | CAMS 176895 | 8,150 | 35 | 9003 | 9265 | 95.45 | Upper Beds | 35 |
| UW 54757 | LLNL | CAMS 186411 | 14,450 | 360 | 16644 | 18630 | 95.4 | Grey Silt | 44 |
| UW 52702 | LLNL | CAMS 172884 | 19,480 | 100 | 23164 | 23775 | 95.45 | Cobbly Grey | 60 |
| UW 51656 | LLNL | CAMS 179248 | 19,480 | 90 | 23185 | 23770 | 95.45 | Upper Red | 60 |
| UW 53734 | LLNL | CAMS 179241 | 20,000 | 100 | 23815 | 24238 | 95.45 | Cobbly Grey | 70 |
| UW 51516 | LLNL | CAMS 170433 | 20,140 | 60 | 23896 | 24412 | 95.45 | Cobbly Grey | 88 |
| UW 52730 | LLNL | CAMS 176886 | 20,270 | 80 | 24120 | 24625 | 95.45 | Cobbly Grey | 88 |
| UW 52713 | LLNL | CAMS 172888 | 20,850 | 120 | 24791 | 25581 | 95.45 | Cobbly Grey | 92 |
| UW 52732 | LLNL | CAMS 176887 | 21,060 | 90 | 25184 | 25662 | 95.45 | Cobbly Grey | 130 |
| UW 52733 | LLNL | CAMS 176888 | 21,510 | 90 | 25703 | 25968 | 95.45 | Cobbly Grey | 130 |
| UW54558 | LLNL | CAMS 186413 | 23720 | 140 | 27641 | 28286 | 95.4 | Mottled Silt | 140 |
| UW54521 | LLNL | CAMS 186412 | 27710 | 220 | 31178 | 32112 | 95.4 | Pebbly Brown | 168 |
| UW 52717 | LLNL | CAMS 176876 | 29,410 | 1,710 | 30862 | 39246 | 95.45 | Pebbly Brown | 175 |
| UW 52721 | LLNL | CAMS 176877 | 30,010 | 260 | 34065 | 35129 | 95.45 | Pebbly Brown | 175 |
| UW 52701 | LLNL | CAMS 172883 | 31,030 | 400 | 34626 | 36192 | 95.45 | Pebbly Brown | 181 |
| UW 52724 | LLNL | CAMS 176881 | 33,500 | 1,250 | 35849 | 41278 | 95.45 | Pebbly Brown | 189 |
| UW 53731 | KCCAMS | CAMS 216524 | 33,560 | 230 | 37597 | 39220 | 95.45 | Pebbly Brown | 200 |
| UW 52722 | LLNL | CAMS 176880 | >29200 | N/A | N/A | N/A | N/A | Pebbly Brown | 200 |

### 4.2 Stratigraphy

There is abundant evidence that water was the primary agent in the deposition of the various stratigraphic units in NTC. The presence of banding, gravel lenses, pronounced erosional surfaces, and the distribution of disarticulated bones all point to the action of running water. Silt and clay constitute the bulk of the cave sediments. Albanese (1976, Unpublished Report) also noted that the majority of sediment (excluding obvious breakdown) is dominated by fine grained clastic material exhibiting sedimentary structures consistent with deposition by moving water.

Sediment flux into the cave from the surrounding surface is likely sourced from the erosion of proximal outcrops of Amsden Formation and soils on the plateau, as well as more broadly sourced aeolian contributions (Stock et al., 2006). Additional sources include breakdown and weathered host rock and paleokarst infill. Multiple surficial processes are likely providing material that is derived from or subsequently blown into the catchment where it can be introduced to the cave by surface water. For instance, aeolian transport of local fine-grained sediment into the immediate catchment can be stored and then mobilized by surface runoff where it is entrained and redeposited below the cave entrance.

Four measured stratigraphic sections demonstrate a relatively consistent pattern of deposition, which can be divided into four fundamental beds that we label the Upper Beds, Primary Beds, Rainbow Beds, and the Lower Beds (listed youngest to oldest). The four beds were further differentiated into 11 identifiable strata that were assigned lithostratigraphic names based on characteristic features (Table 1). An important criterion for determining stratum boundaries was the traceability of a boundary across exposed surfaces.

The Upper Beds and Rainbow Beds are traceable from the South Wall section (Fig. 4A) to the Reference Section (Fig. 4B), which is used to tie all sections together. The Primary Beds are absent in the South Wall section due to erosion prior to the deposition of the Upper Beds that unconformably overlie the Rainbow Beds. This unconformable contact is also traceable to the Reference Section where the Upper Beds overlie the Cobbly Grey. The erosive surface can further be traced to the Saddle Area (Fig. 4C) section where the Grey Silt and Blocky Brown gradually appear as the degree of truncation decreases. This section terminates near the base of the Primary Beds (lowest point of excavation at time of observation). The West Wall (Fig. 4D) section encompasses the Lower Beds through the majority of the Rainbow Beds where it is truncated within the Mottled Silt due to pre-1985 excavations that removed Primary Beds and younger strata.

**Figure 4.**
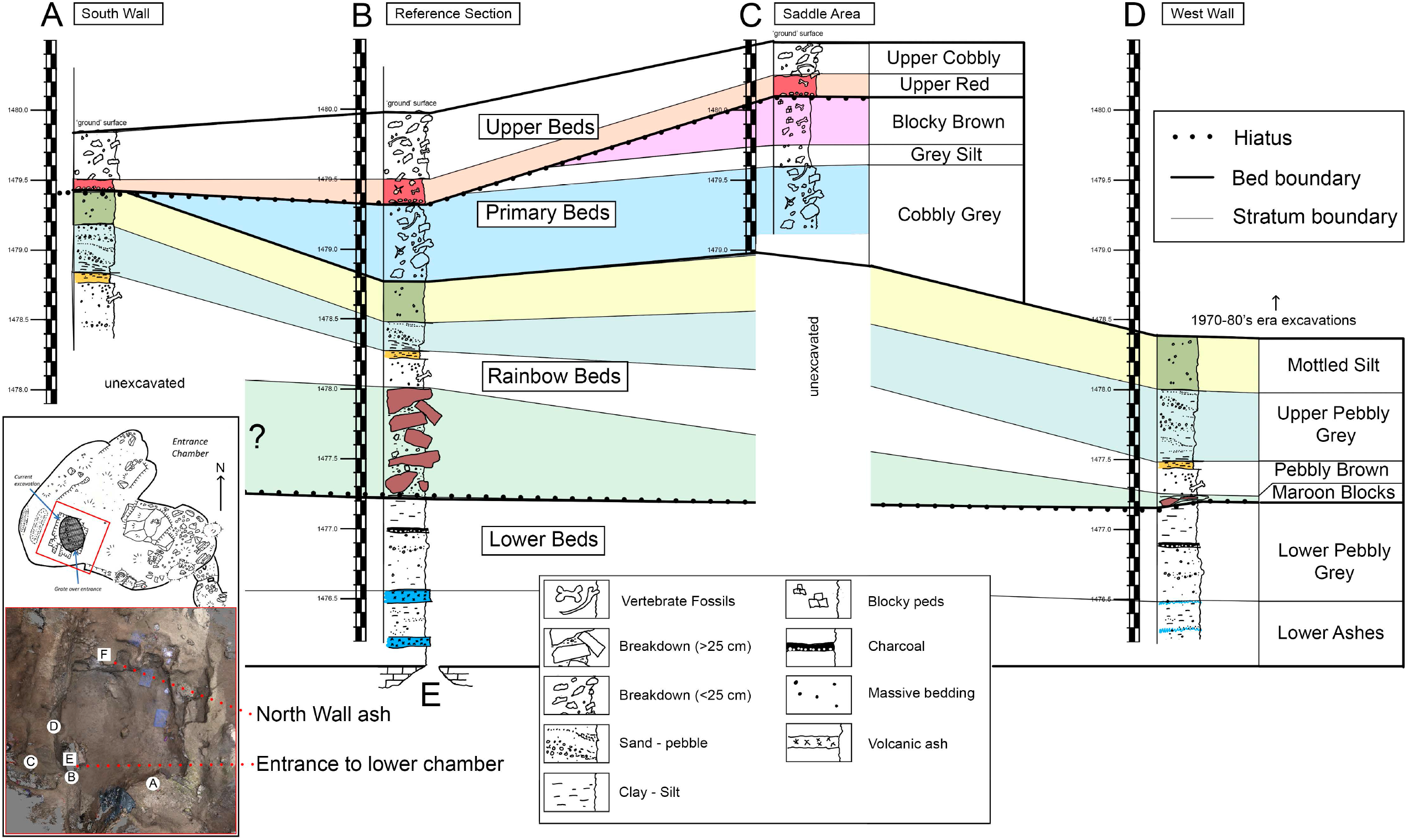
Stratigraphic profiles of four measured sections and their lateral correlations. Inset map (lower left) shows column locations (letters in circles correspond to stratigraphic profiles). Entrance to Lower Chamber (E); North Wall ash (F).

#### 4.2.1 Unit descriptions

The Upper Beds are composed of two intervals, the Upper Cobbly and the Upper Red. Both the Upper Cobbly and Upper Red beds preserve vertebrate remains, but only the Upper Cobbly exhibits abundant pebble to cobble sized breakdown. The Upper Red is a conspicuous red to pale reddish-brown clay-rich silt unit that varies from 2–20 cm in thickness, whose base contains highly-weathered pebble to medium gravel sized breakdown blocks with a chalky appearance (Fig. 5A-C). The lower contact of Upper Red is sharp in the western part of the quarry and irregular but quite apparent along the south wall. This boundary exhibits a notable erosive surface whose incision depth (≥ 1.5 m) increases from the Saddle Area section to the South Wall representing a significant disconformity in both the Reference Section and the South Wall stratigraphy (Fig. 4). Below the boundary, the Blocky Brown is the youngest unit of the Primary Beds, and features well developed fine blocky peds that increase in size from mm to cm scale over ca. 20 cm of depth. Argilans are not observed, though the diffuse and gradational contact with the underlying clay-rich Grey Silt suggests that some degree of illuviation occurred during pedegenisis. Numerous molds of fibrous plant remains were also encountered in both the Blocky Brown and the Grey Silt, though no apparent root traces were documented in either. There is a gradual transition between the Grey Silt and the Cobbly Grey units as breakdown content increases rapidly in both size and abundance. This transition and the upper two units of the Primary Beds are only visible in the Saddle Area, which is effectively the southern edge of the pre-Meachen excavations on the west side of the main quarry. The Upper Beds and Primary Beds make up the vast majority of the earlier excavation; Albanese (1976, Unpublished Report) demonstrated a similar relationship between his Unit 4 (=Upper Beds) and Units 3 and 2 (= Primary Beds).

**Figure 5.**
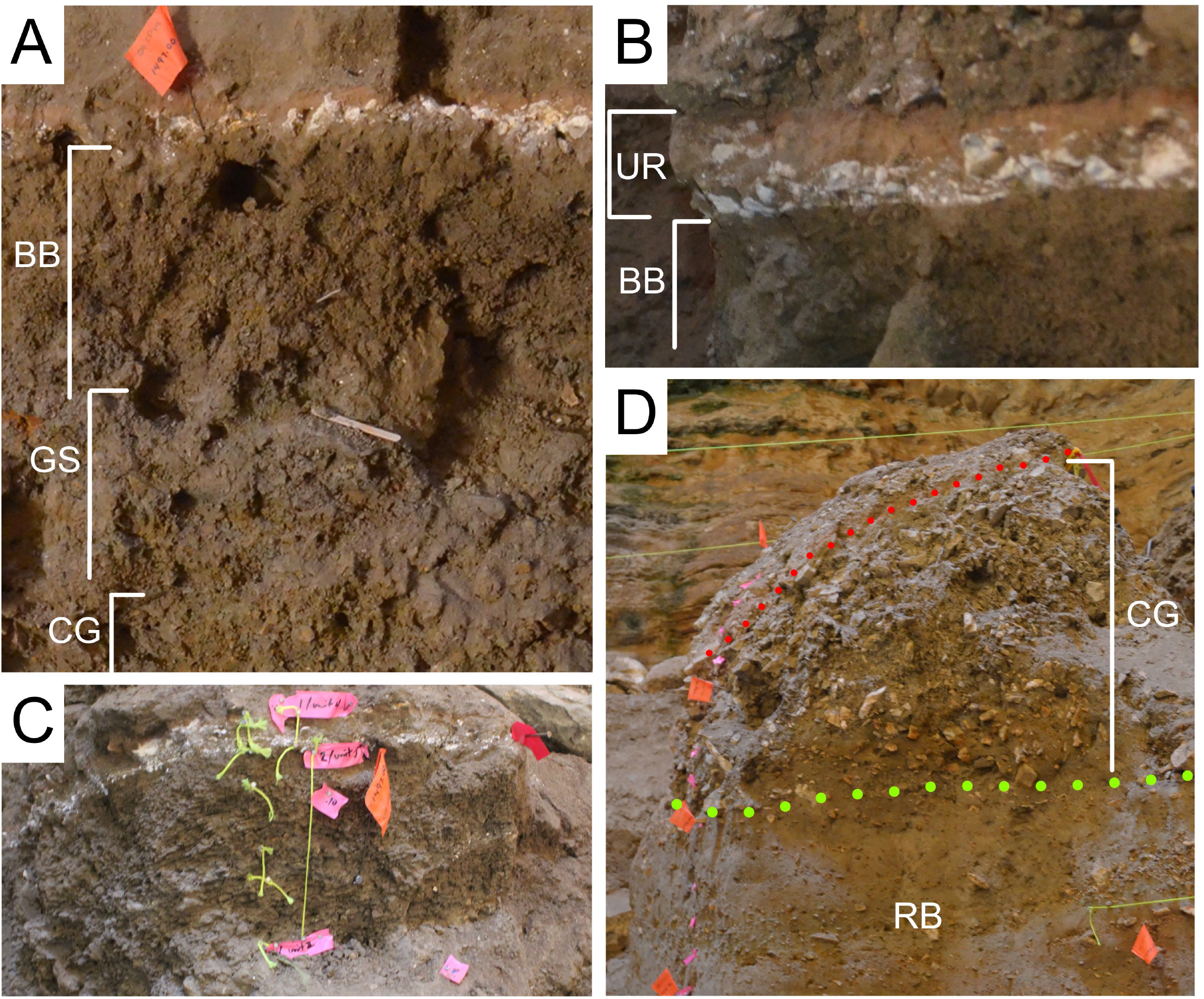
Selected features of the Upper Beds and Primary Beds as seen in the Saddle Area (A-C) and the Reference Section (D). UR = Upper Red; BB = Blocky Brown; GS = Grey Silt; CG = Cobbly Grey; and RB = Rainbow Beds. Red dotted line = contact of UR and CG, note angular incision from right to left. Bright green dotted line = contact between CG and the underlying Rainbow Beds.

The otherwise silt-rich Cobbly Grey is quite apparent because of the high concentration of fine to very coarse gravel, with rare boulder-sized breakdown clasts. It is also noteworthy for the high concentration of megafauna fossils - considerably more than any other unit in the cave to date. The breakdown blocks are angular and show little sign of weathering, sorting, or preferred orientation leading to a massive appearance. The base of the Cobbly Grey is marked by a rather abrupt loss of the gravely component, although the contact is diffuse it does appear to be relatively conformable with the underlying Rainbow Beds (as seen along the southern west wall near the Reference Section, e.g. Fig. 5D).

Relative to the overlying Upper Beds, the Rainbow Beds do not contain nearly as many vertebrate remains (Martin and Gilbert, 1978a; Gilbert and Martin, 1984). Gilbert and Martin (1984) denote the presence of *Panthera* and ‘wolverine’ in the unit below the main bone bearing unit (our Cobbly Grey), but little else. One notable *Panthera* specimen (KU4409) from Albanese’s sub-unit 4 (our Mottled Silt) was dated to 27.8–28.7 cal ka BP (OxA 10078; Williams, 2009). An additional ^14^C determination was recently acquired just prior to submission yielding a 27.6–28.3 cal ka BP (CAMS 186413) date from a specimen of *Vulpes* (UW 54558) collected in the uppermost Rainbow Beds (upper Mottled Silt).

The Rainbow Beds represent four relatively conformable units, three of which display little lateral change in thickness; each unit maintains distinct and mappable characteristics. These four units are traceable across the west wall, and all but the lowermost unit is present on the south wall (Fig. 6). The youngest unit of the Rainbow Beds is the Mottled Silt, a clay-rich silt with apparent, but diffuse, redox features consistent with fluctuations in saturation. There are few randomly distributed breakdown blocks in the pebble to medium gravel size range within this otherwise fine unit. The base appears to be conformable with the underlying Upper Pebbly Grey.

**Figure 6.**
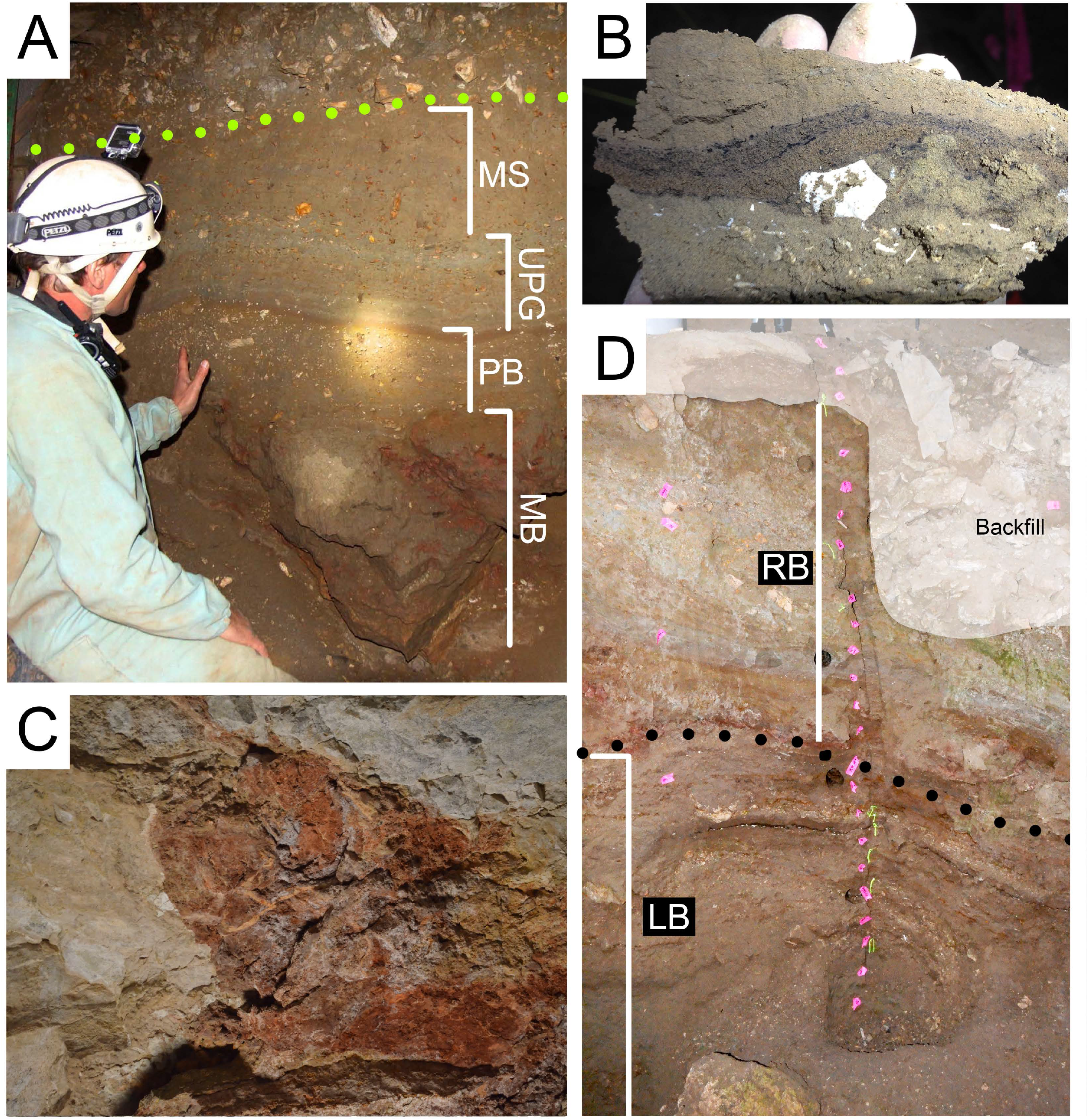
Selected features of the Rainbow Beds and Lower Beds as seen in the Reference Section (A) and West Wall section (B). Dotted green line = Upper Beds/Rainbow Beds contact. Dotted black line = Rainbow Beds/Lower Beds contact. In the upper ¼ of the Lower Beds (B), a laterally continuous charcoal-rich layer is present. C = infilled paleokarst in ceiling lateral to cave entrance. D = charcoal-rich layer (hand sample). MS = Mottled Silt; UPG = Upper Pebbly Grey; PB = Pebbly Brown; MB = Maroon Blocks. RB = Rainbow Beds; LB = Lower Beds.

The top of the Upper Pebbly Grey unit is defined by the abrupt presence of thin (5–20 cm) laterally extensive medium to coarse gravel lenses that show some degree of sorting and lateral accretion suggesting transportation by moving water. These lenses are interbedded with finer grained clay-rich silt, often with mm scale laminations or very low angle cross beds. Distinct color banding is seen in the fine grained intervals throughout this unit. The base of the unit is defined as the top of a paired thin (2–5 cm) but persistent oxidized ‘red’ (10R 4/6) and ‘olive yellow’ (2.5Y 6/6) clay-rich silt layers that abruptly transition to another clay-rich silty unit with abundant pebble to medium gravel breakdown clasts with little to no preferred orientation or sorting giving the Pebbly Brown unit a more massive appearance (Fig. 6A-B). The Pebbly Brown is seen deposited between the upper gaps of boulder-sized breakdown of the underlying Maroon Blocks. The boundary between the Pebbly Brown and Maroon Blocks (Fig. 6B) has only been observed along the west wall; excavations have not extended deep enough to encounter it elsewhere.

The Maroon Blocks are unique within NTC deposits. This unit is most striking in the Reference Section where large (boulder-sized) breakdown with a notable red color, poor sorting, and no preferred orientation are stacked ca. 70 cm thick. Immediately above the Reference Section on the roof of the cave, near the gated entrance, is a portion of ceiling that exhibits the same red blocks that compose (what appears to be) part of brecciated paleokarst infilling (Fig. 6C; Mahan et al., this issue). These blocks become rapidly less common north of the West Wall stratigraphic section where the Maroon Blocks is only a few cm thick and composed of clay-rich silt with the same red color as the isolated large breakdown. This unit, and the overlying units of the Rainbow Beds dips 25-30° N30E below the excavated horizon (e.g., the slump, runoff, and backfilled area of previous excavations where two rungs of an aluminum ladder are exposed, the rest being buried by recent sedimentation) along the northern end of the west wall (Fig. 7). A larger scale unconformity is hypothesized to exist at the base of the Maroon Blocks; regardless, this is the boundary between the Rainbow Beds and Lower Beds.

**Figure 7.**
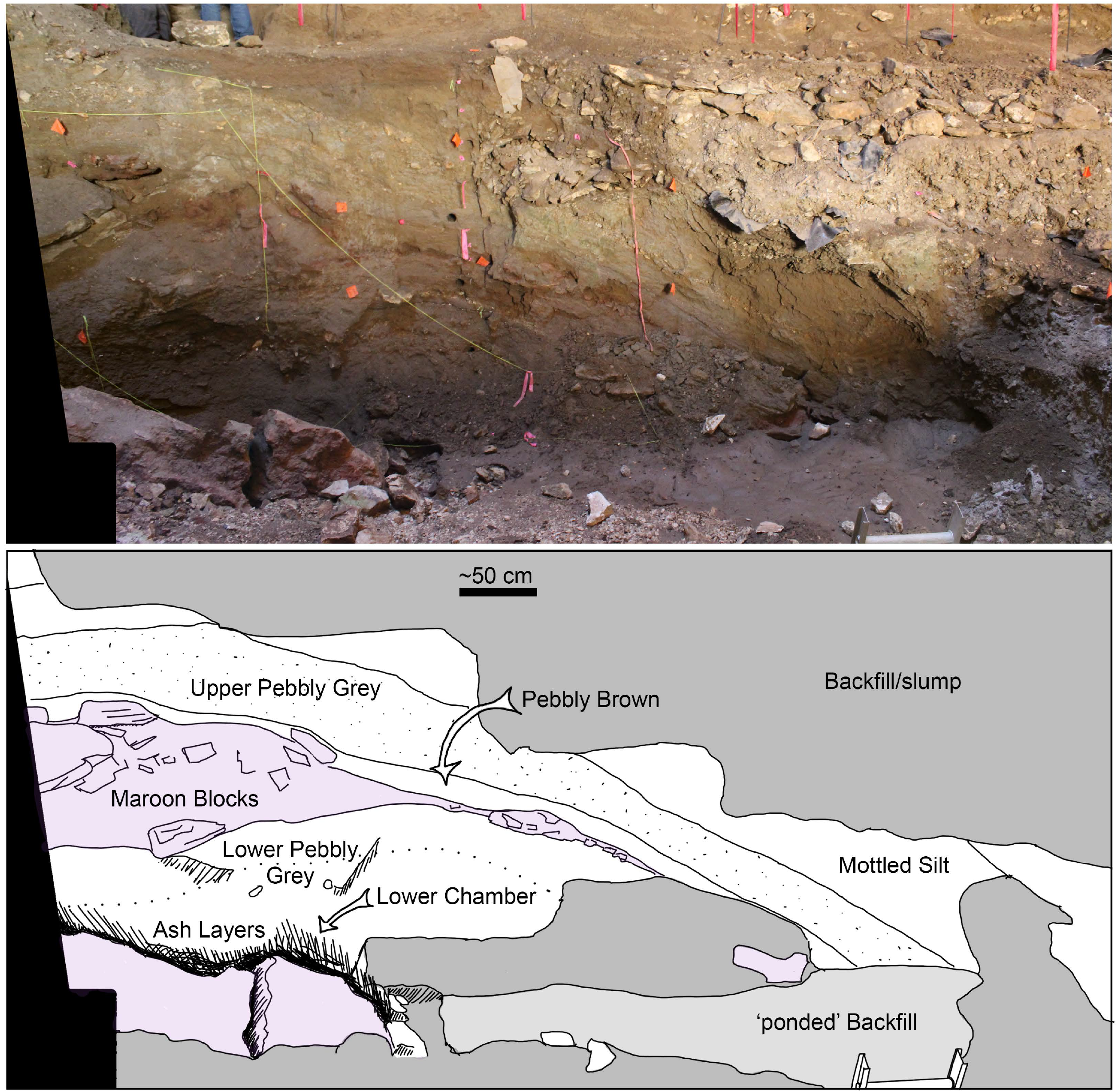
Profile of the west wall (top) with labeled interpretive illustration (bottom). The West Wall section is located vertically below the scale bar (bottom). Note the tip of the ladder buried in sediment accumulated in the ∼2 m deep excavation into the Lower Chamber (ca. 1985; bottom right).

The top of the Lower Pebbly Grey (Lower Beds) sees a return to deposits similar to those observed in the Upper Pebbly Grey, with discrete gravely lenses interbedded with clay-rich silt layers. A 2–5 cm thick charcoal-rich silty layer is a prominent feature of the Lower Pebbly Grey (Fig. 6B,D). The Lower Beds are partially exposed at the base of the West Wall section and they are exposed to bedrock in the Reference Section; however, due to temporary BLM permitting restrictions the Lower Beds were not accessible for more detailed observations at the Reference Section beyond a visual inspection from a distance. The base of the Lower Pebbly Grey is constrained by the occurrence of volcanic ash ca. 40 cm below the laterally extensive charcoal-rich layer.

Two volcanic ashes are present at the base of the Reference Section (Lower Ashes; Fig. 8A-B); these were sampled for geochronology along with an isolated ash (Fig. 8C) on the north wall (Mahan et al., this issue; and Minckley et al., this issue). The ashes are greatly diminished and diffuse at the base of the West Wall section. However, it is apparent that the ash was ultimately deposited by moving water and not via airfall. Sedimentary features demonstrate small-scale low angle bedding (Fig. 8B). Given the limited exposures and lack of access at the time of study, the Lower Beds are the least well understood part of the NTC depositional sequence.

**Figure 8.**
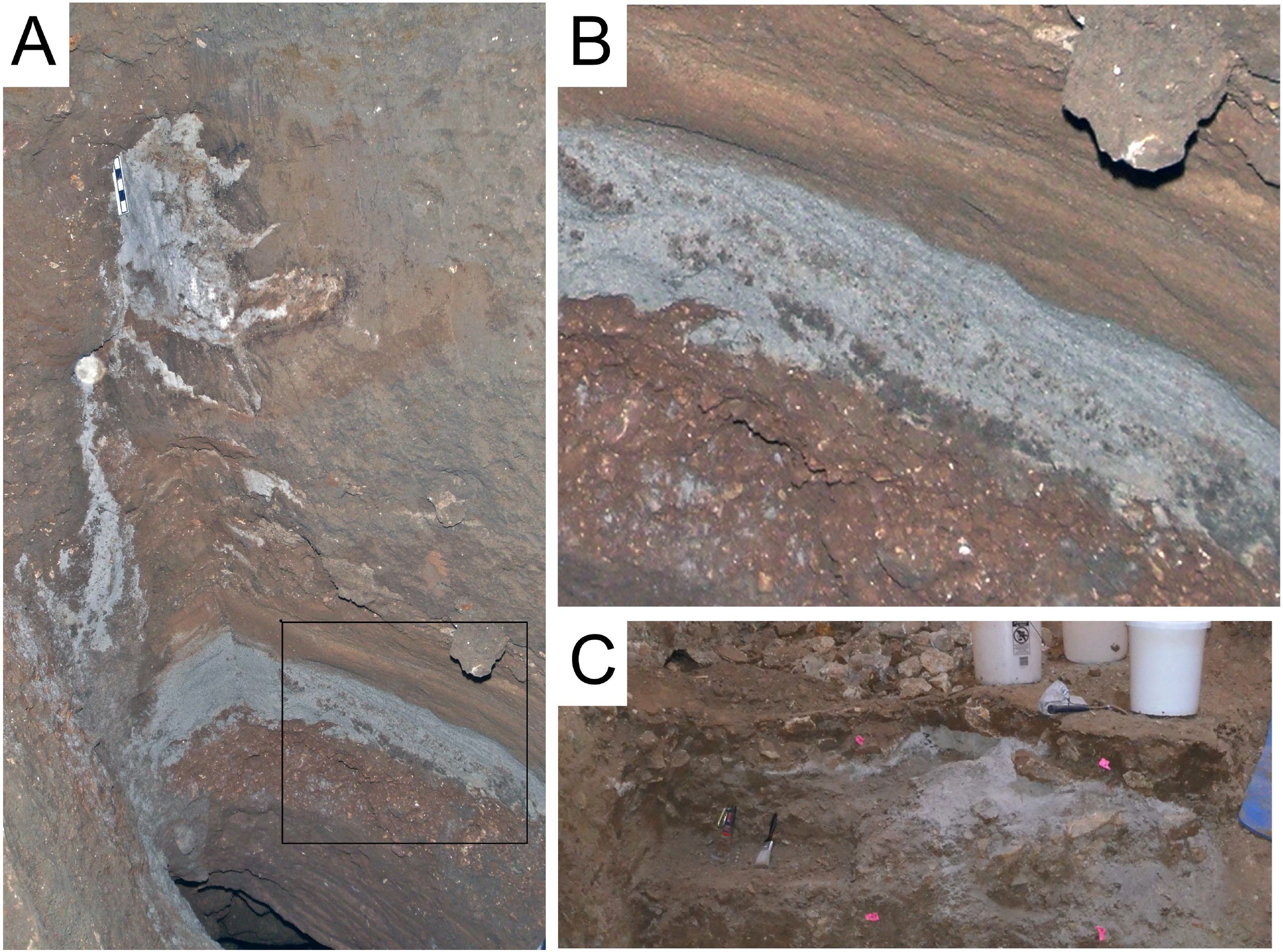
Two grey ash layers (A) were encountered above the entrance to the Lower Chamber (bottom center). The lower ash in (A) exhibits evidence of transport and sorting (B, blowup box in A). Another ash is visible on the north wall (C).

#### 4.2.2 Correlations

The correlation between the Saddle Area, Reference section, and South Wall sections exhibits a significant erosive surface between the Upper Beds and underlying Primary Beds and Rainbow Beds (Fig. 4). The depth of the erosive surface increases with distance from west to east (Saddle Area towards the South wall) along the southern portion of the excavation; this surface can be seen in the reference section where the Blocky Brown and Grey Silt have been completely removed (Fig. 5D; Fig. 9). On the southern end of the east wall the surface dips ∼25°N, where its greatest observed depth extends into the Pebbly Brown (Fig. 9). The remainder of the east side of the excavation (∼495 W line) was covered with excessive backfill or slumped material and fresh outcrop was not encountered for ∼3 m until the 500N line of the excavation (for excavation coordinate system see Meachen et al., this issue) where an unconformity between the Upper Beds and strata that is temporally equivalent to the Primary Beds is seen around 1477.1 m (∼2.5 m below the same contact on the South Wall); this unit was not traceable across the north or east portions of the excavation. Further work is needed to better understand the relationship between this unit and those outlined in this study, but strata that are temporally equivalent to the Primary Beds are present in the northeast corner of the current excavation (500N-515N and 490W-505W) that unconformably underlie Upper Beds that contain a mixture of Holocene and reworked Late Pleistocene vertebrates; this is supported by ^14^C dates.

**Figure 9.**
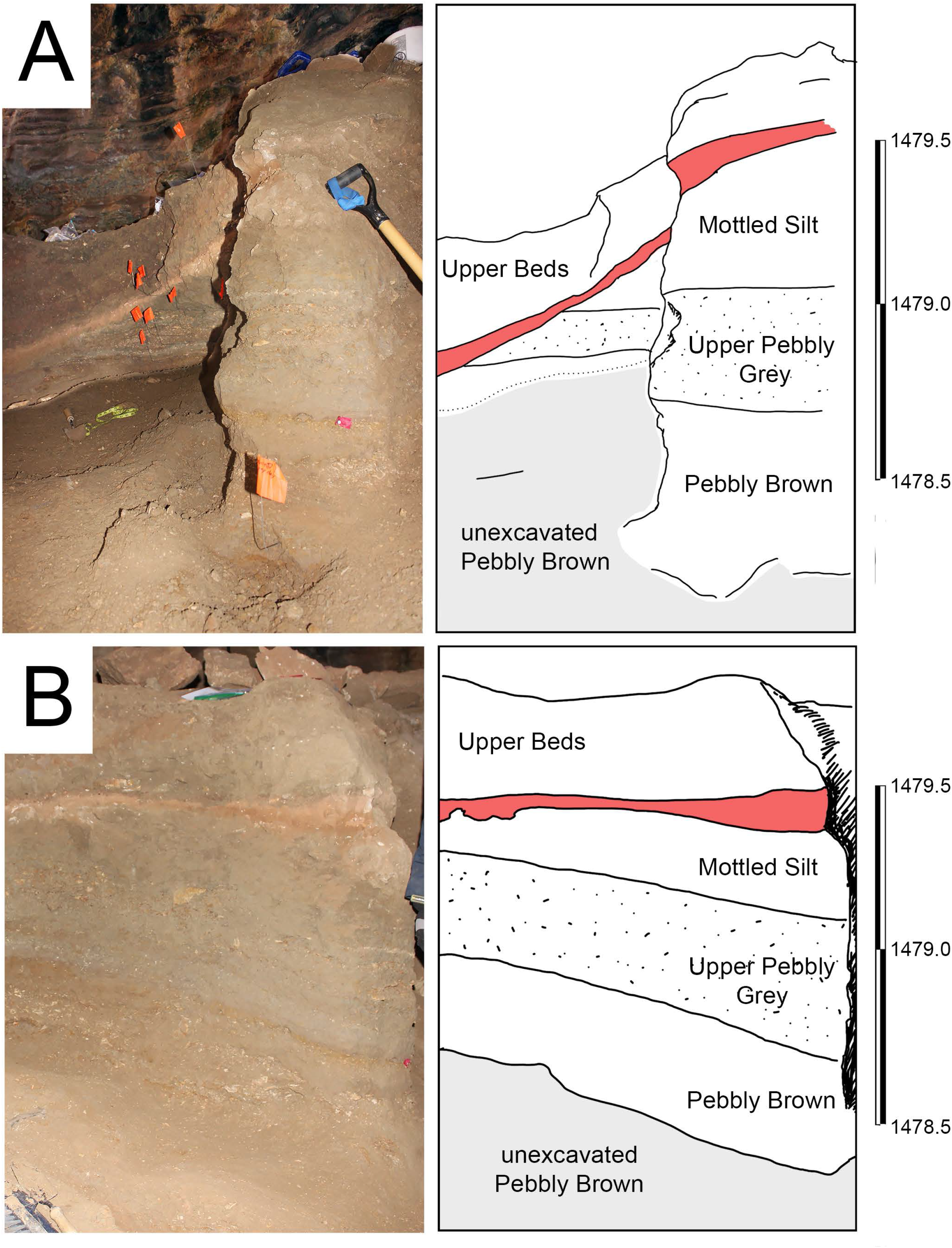
Erosive contact between the Upper Beds and the Rainbow Beds in the southeast corner of the excavation. A = southern East Wall. B = eastern South Wall. Note, in A the contact extends into the Pebbly Brown.

The Reference section is ∼2.5 m south of the West Wall section and units are easily traceable along the exposed surface of the excavation. The strata are northward dipping at ∼30° for the majority of the exposed west wall (Fig. 7). Units of the Lower Beds maintain a relatively uniform thickness and bedding tends to follow the contours of strata below. This is disrupted by the presence of the chaotic assemblage of boulders in the Maroon Blocks, just above the entrance to the Lower Chamber. The Maroon Blocks demonstrate a rapid decrease in thickness north of the entrance to the Lower Chamber. This is linked to a decrease in the size and concentration of breakdown. Excavations from the 1970’s-80’s were backfilled and can be clearly seen in cross section (Fig. 7).

### 4.3 Age-depth model

#### 4.3.1 Radiocarbon ages

Radiocarbon has been the favored approach to NTC geochronology primarily due to the plethora of well-preserved bones in the sediments. Over the course of the last two decades, three different labs have analyzed more than 100 collagen samples extracted from NTC vertebrates to determine the ^14^C age of those specimens (Table S1). Two samples (UW-51516 [OxA 37895; CAMS 170343]; UW-51520 [OxA 37991; CAMS 170433]) were each analyzed at the Oxford Radiocarbon Accelerator Unit (ORAU) and Lawrence Livermore National Laboratory (LLNL) for an interlab comparison. Two other samples (OxA: 14903, 14904 & 14907, 14908) were each replicated at ORAU and another pair (CAMS: 176891, 176897 & 176893, 176898) were each replicated at LLNL for intralab comparisons. Interlab comparisons demonstrate a slightly younger age reported for each LLNL sample relative to the ORAU results. Considering subtle differences in lab protocol and instrumentation, this is not unexpected. Each of the interlab replicates were very consistent with their respective laboratories. Intra- or interlab comparisons with four samples analyzed at Keck Carbon Cycle AMS Facility University of California-Irvine (KCCAMS) were not conducted.

We compiled the results of our analyses and a tabulation of the calibrated ages shows a bimodal distribution (Fig. 10), one that contains post-Pleistocene ages from specimens collected within the Upper Beds, and one that contains the vast majority of total analyzed dates ranging from ca. 23–26 cal ka BP from the Cobbly Grey unit of the Primary Beds. Due to limited exposure of the upper Primary Beds vertebrate material from the Blocky Brown and Grey Silt were inadvertently excluded from ^14^C analyses. The Grey Silt is fully represented in the Saddle Area though it rapidly becomes truncated towards the east and south, while the overlying Blocky Brown is truncated by an erosional surface. We chose one specimen from each unit, both with strong stratigraphic control to further constrain the maximum depositional age of Blocky Brown. Although the specimen from the Blocky Brown did not yield enough collagen for analysis, the Grey Silt specimen (UW 54757) yielded a 16.6–18.6 cal ka BP (CAMS 186411) date, the youngest verifiable Pleistocene ^14^C date for NTC.

**Figure 10.**
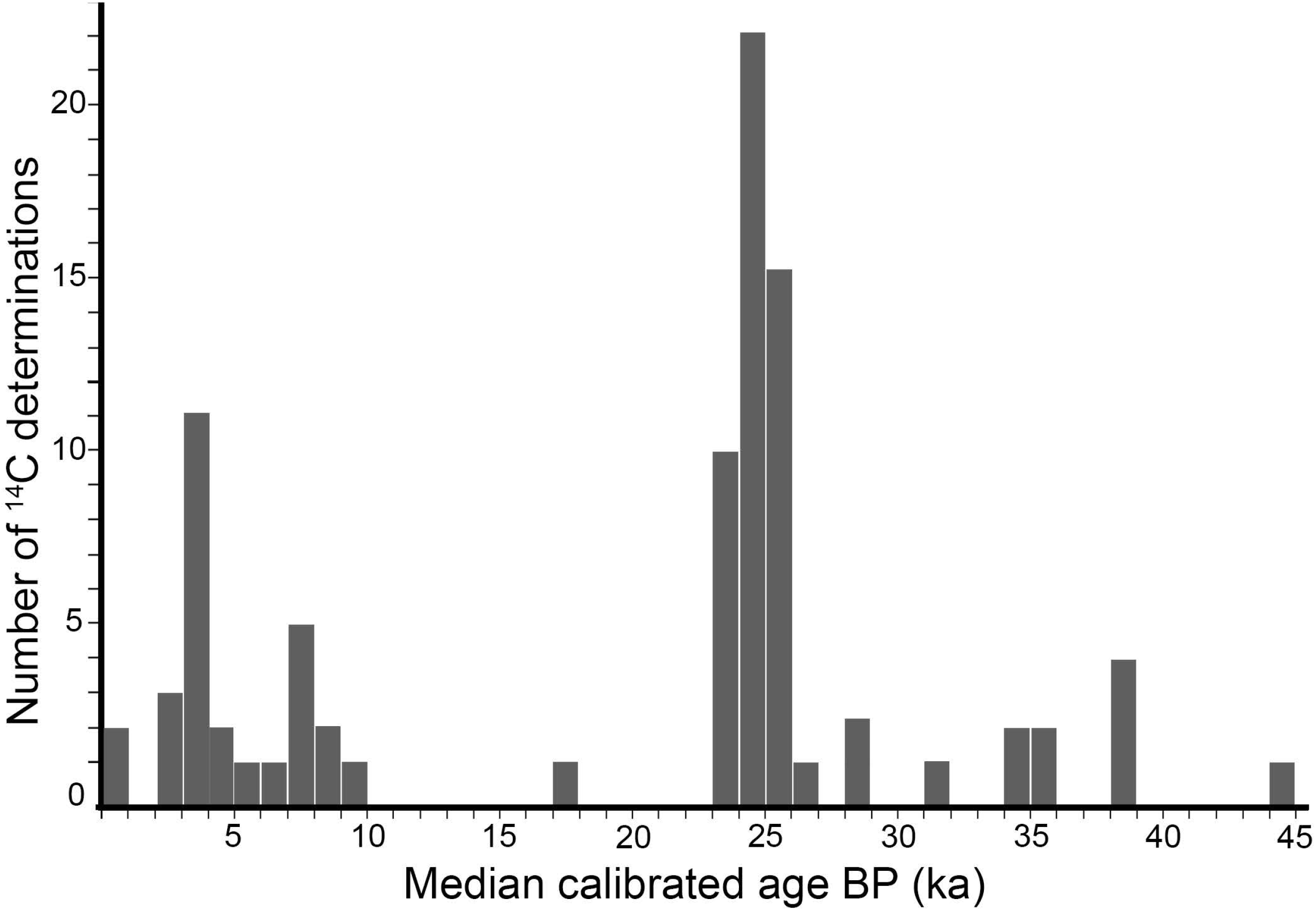
Distribution of median calibrated ages (ka BP) at 95% probability for 97 ^14^C determinations (Table S1); one sample (not plotted) extended beyond the range of the calibration curve.

Three specimens from the Upper Beds have dates that are typical of Cobbly Grey specimens (ca. 23.5, 23.8, & 24.5 cal ka BP [CAM 179248; 179245; 176891]). We suggest that Pleistocene aged specimens collected in the Upper Beds demonstrate reworking of older deposits and are co-mingled with syndepositional post-Pleistocene specimens; therefore, these significantly older specimens are not representative of the age of the deposit. The concentration of vertebrate fossils below the Cobbly Grey appears to be greatly reduced; however, recent excavations have not yet been extended below the Cobbly Grey (e.g., in the Saddle Area, or on the northeast corner), except in specific locations such as the South Wall where Primary Beds are missing. At the South Wall location, the Pebbly Brown is currently the most prolific producer of Pleistocene-aged vertebrate fossils, and these range from ca. 34–39 cal ka BP.

#### 4.3.2 Boundary ages and sedimentation rate

Of the 101 ^14^C age determinations (Table S1), 20 have depth-controlled stratigraphic information, that is to say, their exact position relative to the top and bottom of a particular unit is constrained. The remainder of the ^14^C ages are restricted to unit-level, but not necessarily positionally within their respective units, as the motivation for sampling was for the distributional ages of the faunal remains rather than focused stratigraphic age-depth calculations. The 20 depth-controlled determinations (Table 2) are used as anchors for the age-depth model for the Pebbly Brown and younger units. Two radiometric ages (determined from the lower two ashes) are used to constrain strata below the Pebbly Brown. The median extrapolated age for the base of the Reference Section in NTC is 154.5 cal ka BP, demonstrating that the deposits extend (with recognizable breaks in deposition) from the penultimate glacial period (Bull Lake Glacial) to the Holocene. Three distinct intervals of sedimentation are noted: an upper interval (Upper Beds) that records Holocene accumulation (10.5 cal ka BP to present); a middle interval (Primary Beds through Rainbow Beds) bound by an upper and lower hiatal surface records deposition (53.0 to 17.2 cal ka BP) throughout the Pinedale Glaciation; and a lower interval (154.5 to 132.0 cal ka BP).

**Table 2.** Modeled ^14^C determinations

| Bed | Stratum | Munsell | Description |
| --- | --- | --- | --- |
| **Upper Beds** | **Upper Cobbly** | 10YR 4/4 dark yellowish brown | Silt with weak calcareous cement. Abundant pebble to cobble size limestone breakdown, rare boulder size breakdown present. Abrupt lower contact. |
|  | **Upper Red** | 10R 5/8 red | Prominent silt-rich bed that can be traced laterally across much of the excavated area. Medium to coarse gravel sized limestone fragments with a strongly weathered, chalky appearance. Abrupt unconformable erosive contact with significant incision into underlying units. |
| **Primary Beds** | **Blocky Brown** | 7.5YR 4/4 brown (top)<br>10YR 3/2 very dark greyish brown (base) | Slightly clayey silt with blocky peds that increase in size from fine to medium with increased depth. Clay content increases with depth, and clay skins are present. Fine to medium gravel, diffuse mottles (10R 4/8), and plant fiber molds with oxidized halos are present towards the base. The lower boundary is diffuse. |
|  | **Grey Silt** | 2.5YR 5/3 reddish brown | Slightly gravelly clayey silt with weak calcareous cement. Gradational contact with Cobbly Grey (10–20 cm). |
|  | **Cobbly Grey** | 2.5YR 6/1 reddish grey | Clayey silt with abundant (~20–30% of unit) medium to coarse gravel sized breakdown; boulder size breakdown is rare. Gravel and bone clasts are randomly oriented and poorly sorted throughout the unit. Some diffuse mottling (5YR 5/6; yellowish red) is present. The lower boundary is marked by a distinct abrupt change to a near absence of breakdown. |
| **Rainbow Beds** | **Mottled Silt** | 10YR 6/3 pale brown | Clayey silt with diffuse mottling (5YR 5/8; yellowish red) and rare (<5%) matrix supported medium to coarse gravel. The lower contact is wavy and abrupt. |
|  | **Upper Pebbly Grey** | 7.5YR 4/2 brown | Clayey silt with abundant lenses and accretion sets composed of moderately sorted fine to very coarse gravel at the top of the unit, discrete lenses are more widely distributed within the clayey silt matrix. Lower contact is distinct. |
|  | **Pebbly Brown** | 7.5YR 4/4 brown | A distinct 'purple' (10R 4/6; red) clayey silt layer (2–5 cm thick) marks the upper contact. This is followed by a distinct 'ochre' (2.5Y 6/6 olive yellow) clayey silt layer (2–5 cm). Underlying the 'ochre' layer is a clast rich clayey silt matrix with abundant fine to coarse gravel distributed in shallow lenses. The contact is irregular when following boulder sized clasts of the Maroon Blocks, otherwise abrupt. |
|  | **\* Maroon Blocks** | 10R 4/6 red | Boulder sized breakdown (20–60 cm) is randomly oriented, though it rapidly pinches out to medium to coarse cobbles as the bed dips northward where exposed on the west wall. Silt matrix with randomly oriented fine to coarse gravel size breakdown fills between boulders. Abrupt lower contact. |
| **Lower Beds** | **\* Lower Pebbly Grey** | 7.5YR 4/2 brown | Clayey silt matrix with broadly distributed discrete thin (2–5 cm) lenses of fine to medium gravel. A laterally traceable thin (2–4 cm) charcoal rich layer with fine to coarse gravel sized limestone fragments with a strongly weathered, chalky appearance is present ca. halfway through this interval. |
|  | **\*\* Lower Ashes** | 5YR 6/2 light reddish brown | Clayey silt matrix. Two discrete layers composed of volcaniclastic material are present. The lower ash exhibits sorting and low angle cross bedding. Otherwise similar to Lower Pebbly Grey. |

\* = exposure limited to the west wall
\*\* = exposure limited to entrance to Lower Chamber laterally for ~1 m.

Unlike the clear erosional surface of the upper hiatus (between the Upper Beds and Primary Beds), the lower hiatus (between the Rainbow Beds and Lower Beds) is much less clear. We hypothesize that the base of the Maroon Blocks, which exhibits the most dramatic change in lithology, represents an event that initiates a new interval of deposition after a ca. 80 ka hiatus. We used the basal depth of the Maroon Blocks (∼269 cm) in the Reference Section as the boundary for the second hiatus in the model. Although our model results suggest sedimentation likely ceased somewhere around 132 ka BP, it should be noted that we did not include the 110 ka BP age for an ash that was dated using fission track ([abstract] Gilbert et. al., 1980; Mahan et al., this issue); we have no stratigraphic control for that ash nor was it encountered during the course of our study. Until further dates help to anchor the Maroon Blocks and the Lower Pebbly Grey, we rely on extrapolated ages for this interval. Undoubtedly, this will be refined as new data become available, though we feel the model honors the available data.

In the Reference Section sedimentation initiated around 153 ka with an accumulation rate of around 115 yr/cm. Sediment accumulation ceases around 132 ka at the start of the lower hiatus. The hiatus is terminated with the onset Maroon Block deposition ca. 53 ka. The rate of sedimentation gradually increases from ca. 200 yr/cm to 110 yr/cm throughout the Rainbow Beds. The transition to Primary Beds deposition is marked by higher concentrations of vertebrate remains and medium to coarse gravel and exhibits a much more rapid rate of accumulation (35 yr/cm). There is an approximately 10 fold decrease in accumulation rate (325 yr/cm) between the top of the Cobbly Grey and the truncated surface of the Blocky Brown. There is only one confirmed ^14^C date (16.6–18.6 cal ka BP; CAMS 186411) from a specimen near the top of the Grey Silt. Further sampling is needed in this interval to better understand the duration of the hiatus. Sedimentation resumes above an erosive contact and exhibits a 250 yr/cm rate of deposition, the second slowest in observed NTC deposits.

The above sedimentation rates and median extrapolated ages for major stratigraphic boundaries are based on 20 well constrained ^14^C ages and two other radiometric ages. Hamilton and Krus (2018) suggest the inclusion of all radiometric dates tied to a study area unless there are reasons for their exclusion (i.e., poor sample prep/yield/lack of historical documentation), and suggest the inclusion of DICARB dates even in light of their inherent issues (*sensu* Reuther and Gerlach, 2005). After randomizing ages constrained to particular units (bound depth ranges), we repeated the earlier analyses with an additional 70 ^14^C ages (Table S1) including DICARB 1687. The results were quite similar, with the exception of the 40 cm above the upper hiatus where the larger dataset hypothesizes two intervals of rapid followed by slow rates of accumulation rather than relatively stable rate of accumulation (Fig. 11). Tighter stratigraphic control of Holocene samples will help resolve this issue with additional dating in this part of the section.

**Figure 11.**
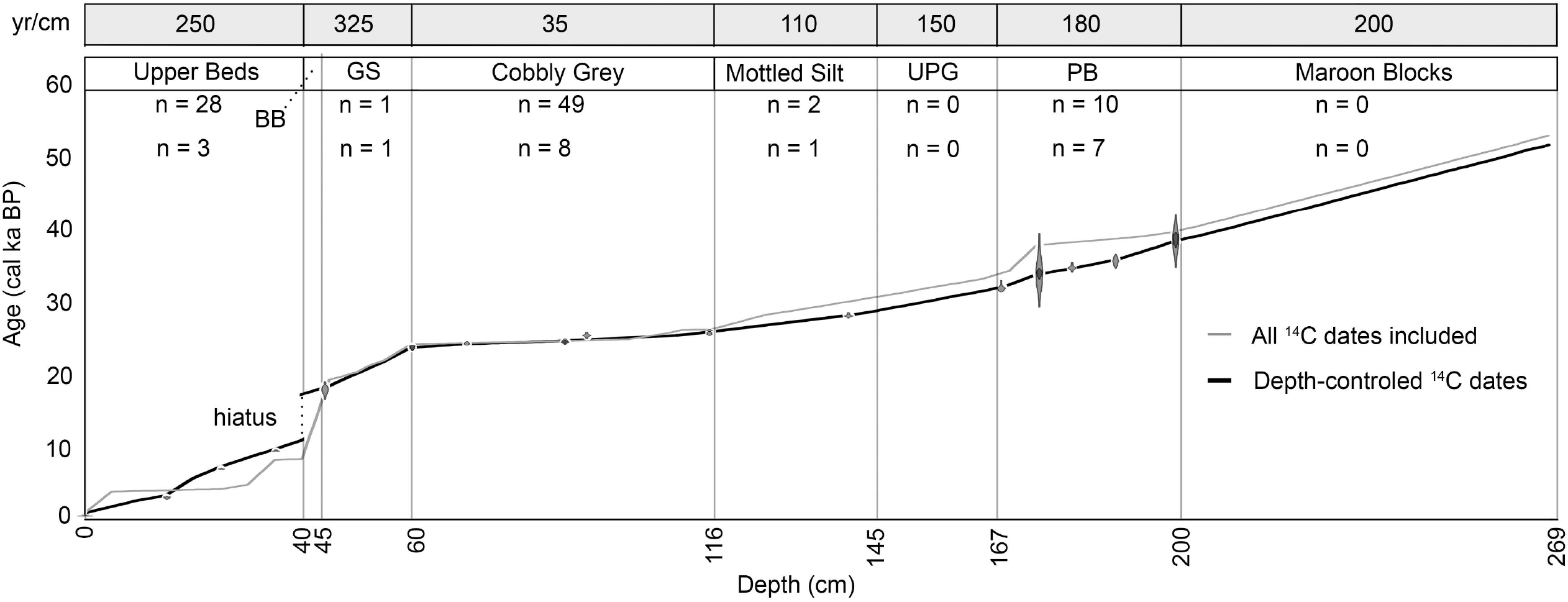
Results of the age-depth model and rates of sedimentation (yr/cm) for the first 50 ka. Analyses with stratigraphically well constrained samples (n=20; black line; data points are plotted with error) and all ^14^C determinations (n=101; orange line) are plotted for comparison. No samples were analyzed from the Blocky Brown (BB). GS = Grey Silt; UPG = Upper Pebbly Grey; PB = Pebbly Brown.

## 5. Discussion

Reconstructing the temporal relationships between sediment (and bone) accumulation, paleoenvironment, and paleoecological circumstance is critical to understanding the context and history of paleontological localities such as NTC. The fossil-rich deposits have proven to be a treasure trove of vertebrate stories ranging from the preservation of genetic material, trophic associations, paleoecology, paleogeography (e.g., Mitchell et al., this issue; Minkley et al, this issue; Higgins et al., this issue; Redman et al., this issue). However, outside of the unpublished work by John Albanese (1976), little has been done to further our understanding of stratigraphic relationships of excavated sediments. Moreover, confusion has arisen as various authors have created their own nomenclature to apply to NTC deposits without the requisite correlations needed to maintain continuity across studies (e.g., Martin and Gilbert, 1978a; Gilbert and Martin, 1984; Wang and Martin, 1993; Williams, 2009).

### 5.1 Stratigraphic Nomenclature

A hand-drawn note (Appendix, Fig. S1) found amongst NTC fieldnotes and files curated at the University of Kansas Biodiversity Institute Division of Vertebrate Paleontology is the only known correlation of the ‘zone’ and ‘unit’ systems of stratigraphic nomenclature created in 1976/1977: the ‘zone’ system was first devised by SAC (S.A. Chomko - 1976) and modified by SAC/KAL (S.A. Chomko & K.A. Lippencott - 1977) and Albanese’s ‘unit’ system of nomenclature (Albanese, 1976 Unpublished Report). Unfortunately the long-term utility of the latter was complicated by the inverted nature of the original unit designations. In his 1976 report, Albanese described 11 discrete stratigraphic intervals (numerically descending with stratigraphic depth). He later (1977) amended his original report by adding hand-written notes assigning each of his 11 units to one of four primary ‘Units’ ([dataset] Lovelace et al., 2021). In other words, Unit 1: sub-unit 1 was the oldest stratigraphic unit known at the time of his description, and Unit 4: sub-unit 11 was stratigraphically youngest. The ‘zone’ nomenclature along with Albanese’s ‘unit’ nomenclature were used extensively during the 1974-1985 excavations to record the stratigraphic context of fossil specimens. Issues arose when excavations began to encounter stratigraphic layers underlying Albanese’s Unit 1: sub-unit 1.

It is unclear when a shift in nomenclature occurred. The first example is found in Martin and Gilbert’s (1978a, fig 5) schematic stratigraphy figured with fauna typical of each unit along with an abbreviated description of seven stratigraphic units; they labeled their units Stratum 1–7 (youngest to oldest, respectively). While there is utility in this nomenclatural shift (e.g., it is easier to add new units as they are discovered below the last labeled unit), the new stratum intervals do not correlate 1:1 with the ‘zone’ or ‘unit’ system previously established, and the published strata lacked detailed descriptions or correlations with Albanese’s prior descriptions. In addition, descending numerical stratigraphic nomenclatures can potentially complicate issues as new strata may be discovered between existing units. Six years later, Gilbert and Martin (1984, fig. 6.3) published a simplified stratigraphic diagram (which appears to be modified after Albanese’s A-A’ cross section; Albanese, 1976 Unpublished Report) to demonstrate the relative position of three ashes (found in 1979) interbedded with cave sediments found well below Stratum 7 of Martin and Gilbert (1978a). However, none of the stratigraphic intervals were described or labeled and the schematic is inconsistent with their previous publication making a direct comparison between the two difficult.

At least two units have commonality between all stratigraphic schemes (based on unit descriptions/figures), the laterally continuous clay-rich silt with weathered (chalky appearance) pebble to gravel size limestone clasts at the base (our Upper Red), and the main bone bearing unit (our Cobbly Grey) with abundant fine to very coarse gravel breakdown in a grey/green silt matrix (Albanese, 1976 Unpublished Report; Martin and Gilbert, 1978a). Descriptions of an associated *Panthera* specimen (skull = KU 31417; Martin and Gilbert, 1978b) from the just below the contact with the breakdown-rich ‘main bone layer’, (i.e., Unit 2 -sub-unit 5 and Unit 1 -sub-unit 4), allows correlation with Stratum 4-5 of Martin and Gilbert (1978a) and our Mottled Silt unit. Albanese’s Unit 1 (sub-unit 1) is his lowest described stratum and is noted as a clayey light tan silt with abundant angular limestone fragments (fine to very coarse gravel) that occur scattered throughout, and as lenses (Albanese, 1976 Unpublished Report). This is comparable to the Martin and Gilbert (1978a) Stratum 7 and our observations of the Pebbly Brown (see section 4.2.1).

Wang and Martin (1993) used Albanese’s unpublished cross sections and sub-unit designations (not cited), but misidentified the ‘main bone bearing layer’ as Unit 4. They clearly were discussing sub-unit 5, which is the main bone-bearing unit of Albanese, and is even drawn and labeled correctly in their reconstructed fence diagram (Wang and Martin, 1993 fig. 5). We have collated all significant contributions to NTC stratigraphic nomenclature and correlated them with our proposed lithostratigraphic nomenclature, including our extrapolated ages for each unit boundary (Fig. 12).

**Figure 12.**
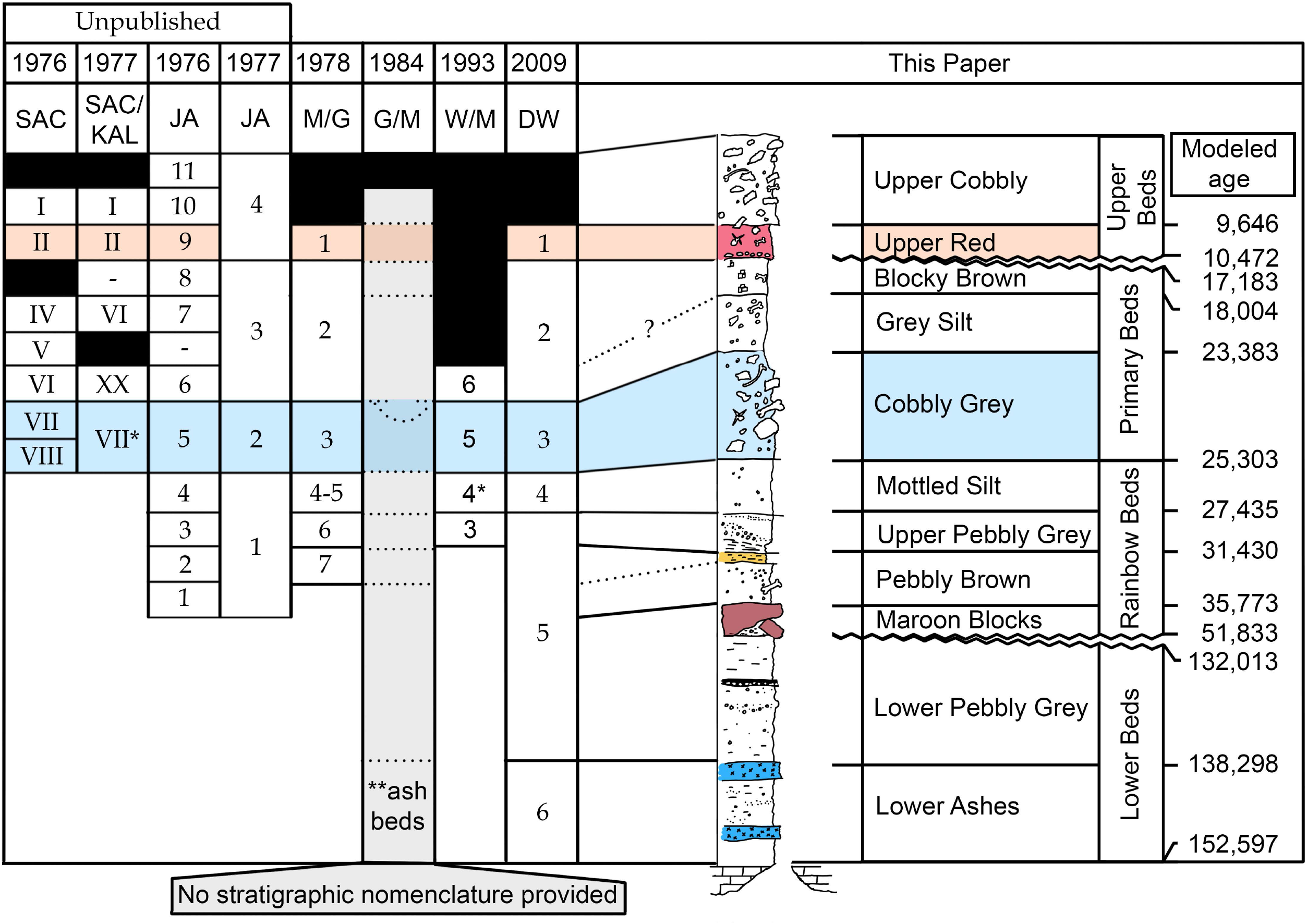
Correlation of historical nomenclature including unpublished and published accounts. The Upper Red (highlighted with light red correlation) and the Cobbly Grey (highlighted with light blue correlation) share commonalities across multiple publications. *Note that Wang and Martin (1993) misidentify Unit 4 as equivalent to ‘zone’ VII and referred to Unit 4 as the main bone-bearing unit; their description and fence diagram (their Fig. 5) is based on sub-unit 5 of Albanese (1976 Unpublished Report). **Inferred position of ashes from Gilbert and Martin (1984).

### 5.2 The ‘snow cone’ hypothesis and the ‘Lower Chamber’

While we cannot fully reconstruct the evolution of deposits within the cave, specific observations do provide insight and allow for the support or rejection of other hypotheses. The possible presence of a year-round snow cone during the Pleistocene deposition was used to explain the displacement and concentration of fossils several meters lateral to the footprint of the cave entrance. The snow cone hypothesis was also invoked to explain unusual topography of the cave floor, namely that the most fossiliferous strata were found not only lateral to (max = <10 m), but also elevated above (≤2 m) the footprint of the cave entrance (Martin and Gilbert, 1978a; Wang and Martin, 1993).

The current average temperature is reported to be around 5° C (Martin and Gilbert, 1978a; Wang and Martin, 1993; Meachen et al., 2016). While modern average cave temperature alone does not preclude the possibility of year round snow during peak glacial periods, it demonstrates that under recent (i.e., interglacial) conditions a permanent snow cone does not exist. It should be noted that permanent snow does occur in several caves in the Cottonwood Creek area south of NTC, and northwest of the cave in the Pryor Mountains; however, those caves are greater than 800 m higher in elevation and are located in more forested areas (Elliot, 1963; Hill, 1976; Higham and Palmer, 2018). Further observations of these caves could be used to compare the resultant sediment accumulation and distribution in the waning presence of a permanent or semi-perminant snow cone. In addition, no periglacial features have been observed in NTC (Albanese, 1976 Unpublished Report). The topographic depression found immediately below the cave entrance was proposed to represent the location of the snow cone, and as a result sediment, bones, and animals, that fell into the cave were proposed to have effectively been shunted laterally to build up around the margin of the cone causing a high concentration of vertebrate remains (Wang and Martin, 1993).

If, as it has long been presumed, vertebrate remains are sourced from animals that accidentally fell into the cave, it must be at least noted that there is the distinct possibility that those animals were in motion (walking or running across the landscape). Under those circumstances, moving animals will maintain the velocity at which they lose contact with the ground for the duration of the fall. Conservatively the vertical descent is 24 m, thus a free fall of this distance would take ∼2.2 s (impact velocity ∼22 m/s). Objects entering free fall with a horizontal velocity less than 1 m/s would fall within 2 m of the footprint of the entrance. Megafauna trotting or running down the ridgeline and entering the cave at higher velocities (i.e., 1.5 to 4 m/s; Higgins et al., this issue) would impact somewhere between 3–9 m from the entrance footprint; momentum could potentially carry their carcasses even farther after impact. If the animal did not rebound off of the opposite side of the cave entrance, it is possible to fall via projectile motion > 20 m horizontal to the fall line. The higher accumulation of vertebrate remains west of and lateral to the cave entrance might well be explained with projectile motion rather than sliding down a snow cone.

We support an alternate explanation for the depression underneath the entrance: that erosion was the major cause of surface deflation observed during earlier investigations. Erosion as a potential mechanism was originally proposed and simultaneously dismissed as unlikely because of a presumed lack of slope capable of allowing material to move laterally (Martin and Gilbert, 1978a) and it was thought that small scale cut and fill structures suggested erosion wasn’t a prominent surface process (Wang and Martin, 1993). However, active redistribution of sediment by moving water has been documented throughout the history of excavation. For instance, curated BLM reports discuss means of controlling erosion between field seasons with plywood structures ([dataset] Lovelace et al., 2021), and observed stratigraphic relationships show clear evidence of erosion as a significant form of surficial modification.

Our stratigraphic sections demonstrate that the Rainbow Beds are at their maximum elevation just below the cave entrance (Fig. 4). Given the discovery of *in situ* vertebrate remains that yielded ^14^C dates from the Mottled Silt (27.8 to 28.7 cal ka BP; OxA-10078) and Pebbly Brown (37.6 to 39.2 cal ka BP; CAMS 216524) under the entrance to the cave during the buildup and into the peak of the Last Glacial Maximum, 21,000 cal yr BP (Pinedale Glaciation; Pierce, 2003; Pierce et al., 2018), the presence of a permanent snow cone is rejected. Late Pleistocene strata equivalent to the Primary Beds most likely existed below the entrance, dipping to the north and west where it was possible for material to be moved laterally downslope. We suggest early Holocene erosion prior to deposition of the Upper Beds is the most parsimonious solution to the missing strata.

Although we propose that Primary Beds equivalent strata under the cave entrance were removed by erosion, it raises the question: where could eroded sediment/bones have gone? There is support for reworking of vertebrate material derived from Cobbly Grey sediments in the Upper Beds (see section 4.2.2), and it is possible that the eroded material was transported down the main talus slope mixing with Holocene vertebrate remains; this may be the case for several bones attributed to Pleistocene-aged mammals found on the surface of the slope in early reconnaissance (Rushin, 1973). Numerous ^14^C age determinations with stratigraphic control will be required to further interrogate this hypothesis in future excavations. The presence of the Lower Chamber is another possible route of sediment transport and deposition of reworked material.

The entrance to the Lower Chamber was not encountered prior to the 1985 excavation indicating that the entrance was covered by breakdown prior to that date. An initial investigation of the Lower Chamber in 1985 ended with the collection of two Pleistocene aged taxa, including a mammoth femur (KUVP 85000) and an equid skull (KUVP 84947; Wang and Martin, 1993). This demonstrates communication with the Upper Chamber at some point with an opening big enough to accept boulder-sized Pleistocene material. Outside of the LiDAR survey of the physical dimensions and limited surface collection of vertebrate remains (see section 4.1) in 2014/2016, little is known about the Lower Chamber and its relationship with the main chamber. It is proposed that the Lower Chamber, at least periodically, acted as a repository for material derived from deposits within the Main Chamber. The complexities of sedimentation in the Lower Chamber may be exacerbated by intermittent sediment supply and will likely exhibit significant temporal mixing of material (including reverse stacking with older on younger material). Understanding the relationship between the Lower and Main chambers is critical to piecing together the history of sedimentation within the cave; this should be a focus of future work in the cave.

### 5.3 Constraining depositional history

Considering the concentrated distribution of vertebrate remains within NTC deposits, we employed the age-depth model to help frame fossil-dense strata (i.e., Cobbly Grey) within a spatial and temporal context. Although there are three hiatus bound sedimentary packages with evidence of truncated or missing stratigraphic units, sedimentation is rather consistent between comparable units (i.e., the Mottled Silt/Upper Pebbly Grey and the Lower Beds) with an ∼110–150 yr/cm accumulation rate; the exceptions are the Cobbly Grey with a much higher rate of sedimentation (∼25 yr/cm) and the upper Primary Beds (>300 yr/cm). In addition, units that are bound above and below hiatal surfaces have comparable lithology, breakdown distribution, clast size, and intra-unit thicknesses (i.e., Upper Pebbly Grey and Lower Pebbly Grey), suggesting similar depositional circumstances. The taphonomic signal is also suggestive of a consistent depositional setting (Redman et al., this issue), though it is unclear why there is an increase not only in depositional rates, but also the dramatic increase in vertebrate remains in the Cobbly Grey.

By demonstrating that stratigraphic boundaries can be traced between sections we suggest that extrapolated boundary ages can also be translated across these surfaces making temporal correlations possible. Furthermore, the extrapolated boundary ages allows us to place fossiliferous intervals into a chronostratigraphic framework (Fig. 13). When compared to local and regional glacial history (e.g., Salisbury and Blackwelder, 1903; Nelson, 1977; Pierce et al., 2018; Minckley et al., 2021), the vast majority of preserved sediments were deposited during the buildup of both the penultimate and Last Glacial maxima. Interglacial deposits were not encountered between the two glacial maxima, though erosion, bypass, and deposition are clearly active processes in the Holocene.

**Figure 13.**
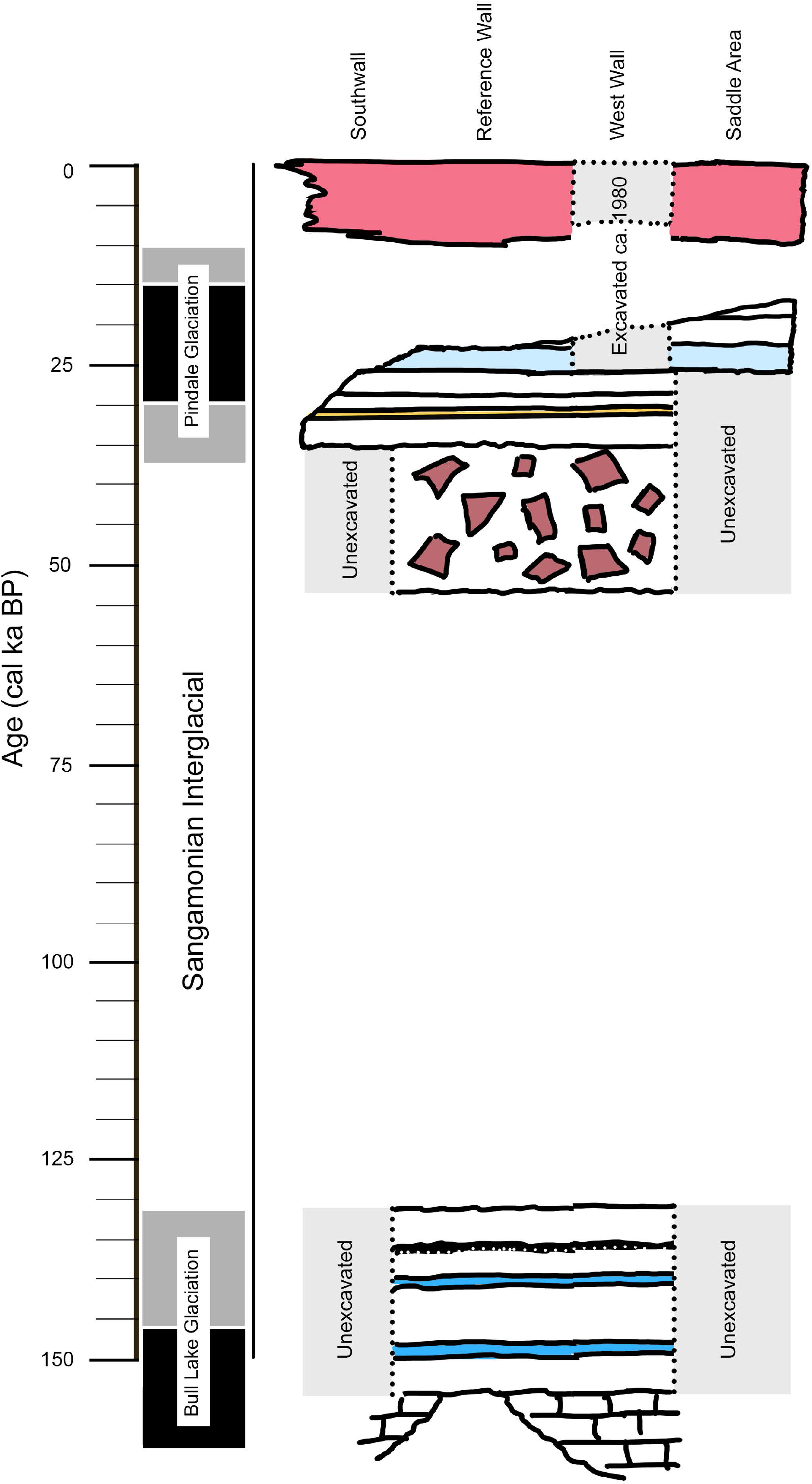
Chronostratigraphic representation of Natural Trap Cave sediment accumulation compared to regional glaciations. Note the lack of preserved stratigraphy deposited during the Sangamonian Interglacial in the area of the main excavation(s). Shaded areas on the left of the timeline represent interglacials (white), buildup/decline (grey), and peak (black) glacial periods. Red = Upper Beds; light blue = Cobbly Grey; dark red = Maroon Blocks; dark blue = lower two ashes.

The base of the Lower Beds in the Reference Section represents the start of sediment accumulation in NTC which coincides with buildup of the penultimate glacial period (Bull Lake Glaciation ca. 150–130 ka; Pierce et al., 2018). The upper surface of the Lower Beds marks the base of the lower hiatal boundary. This surface does not exhibit significant erosion supporting a period of bypass rather than downcutting, at least in the exposures of the Reference Section. The hiatus is terminated with the onset of deposition at the base of the Maroon Blocks which has an extrapolated age of 53 cal ka BP. The Maroon Blocks represent a significant event composed of large red-colored breakdown blocks. The thickest accumulation is observed at the base of the Reference section and on both the east and west side of the entrance to the Lower Chamber. A conspicuous portion of the cave roof immediately over the Reference Section appears to be dark red paleokarst infill that may have been the source of the breakdown (Fig. 6C). We tentatively hypothesize that this event resulted in a physical barrier that altered the topography of the floor, or plugged the entrance to the Lower Chamber, reinitiating deposition in the main excavation area throughout the Pinedale Glaciation.

Sometime near the end of the Pleistocene, erosion likely began to reshape previous deposits resulting in the mixing of Holocene and Pleistocene aged faunas in the Upper Beds. The record of Early Holocene through present is more consistent with deposition of eroded Pleistocene material along with more contemporaneous specimens and sediment. Presently erosion and bypass are dominant processes within the cave as material is transported to the lowest portions of the excavation (which is currently bowl-shaped) and overflow into the Lower Chamber. Given the active surficial processes seen in the cave today, and depositional history throughout the last ca. 150 ka, it seems unlikely that sediment input from the surface would be halted for >70 ka. At minimum, aeolian contributions would be expected. It is more likely that warmer and wetter interglacials were (are) relative periods of erosion and bypass rather than deposition in the area of the main excavation.

A final point to note: the Saddle Area is our only measured section that contains the Grey Silt and Blocky Brown, which correlate to Albanese’s Unit 3 (sub-units 6–8). Given the increasing presence of the Blocky Brown to the east as the depth of the erosive unconformity lessens in that direction, it is possible that the Blocky Brown west and north of the Saddle Area (i.e., Martin-Gilbert excavations) were thicker than in the section we observed. This would effectively reduce the amount of time missing between the Upper Red and underlying Blocky Brown. For this study, we have discounted most of the early DiCARB data because of the inclusion of multiple specimens (to get enough collagen to analyse at that time) and/or the original analytical data are not available. However, three of the DiCARB dates fall *within* our modeled upper hiatus (10.9, 12.8, 14.7 uncalibrated ^14^C ka BP). Our final ^14^C analysis (in 2021) yields one date from the upper Grey Silt at the older end of this range (i.e., 14.5 uncalibrated ^14^C ka BP; CAMS 186411). It is possible that the DiCARB dates reflect reasonable radiocarbon ages, but were not reproduced in our studies because the strata they derived from were absent (eroded) from the sections we studied. A systematic study to determine ^14^C dates from specimens collected during Martin-Gilbert excavations within a single square with known depth and stratigraphic position spanning sub-units 5 through 11 would help establish if the erosional surface extends to this portion of the excavation, or if the Pleistocene-Holocene boundary is preserved as hypothesized by previous workers (Martin and Gilbert, 1978a; Gilbert and Martin, 1984). If the Pleistocene-Holocene boundary is found to exist in NTC, further study or continued excavation in the western and north of the deposits is warranted considering the paucity of sites that record this important transitional period.

## 6. Conclusions

- Using lithostratigraphic terminology we have established an easily modifiable system to incorporate new units as they are discovered, regardless of their position in the profile relative to existing units. This new system is correlated with previous stratigraphic nomenclatures to establish a consistent framework for future excavations and to allow the comparison of current and future work with previously published results that form the backbone of our understanding of NTC.
- We used more than 100 radioisotopic dates to develop an age-depth model that enables us to assign an extrapolated age to each stratigraphic boundary. These boundaries are traceable between sections allowing us to place deposits throughout NTC excavations into a chronostratigraphic framework making broader correlations and comparisons possible.
- Counter to the original hypothesis that NTC represents continuous deposition over the last 110 ka, we demonstrate that deposition likely began around 155 ka and that there is an ∼75 ka period of non-deposition in the study area between the penultimate and Last Glacial maxima.
- There appears to be a relationship between sediment accumulation in the main excavation area and the buildup through peak glacial periods. Interglacial periods are not represented in the stratigraphic assemblage and may reflect a period of erosion, bypass, or nondeposition. Warmer and wetter conditions of the interglacials would likely see more influx of water, and sediment may have been transported further downslope or into the Lower Chamber (e.g., bypassing the main excavation area).
- Latest Pleistocene to earliest Holocene strata are not present in the four stratigraphic sections measured for this study. It is possible that strata of this age were removed during early Holocene erosion prior to onset of deposition that preserves a mixture of Holocene and reworked Late Pleistocene vertebrate remains. Additionally, it may be possible that the erosional surface is restricted to the southern and eastern portions of the excavation (our study sections), allowing for the potential preservation of the Pleistocene-Holocene boundary in historically excavated (and yet to be excavated) areas in the northwestern areas of NTC deposits.

## Author contributions

Conceptualization, DML, CMR, TAM; Methodology, DML, TAM; Investigation, DML, CMR, BWS, JL; Formal analysis, DML, TAM; Data Curation, DML, CMR, KB; Resources (^14^C data), KJM, AC, LF, HH; Resources (cave safety), JL; Visualization and Writing (Original draft), DML; Writing (Review and editing) all authors; Funding acquisition, JLM, JAM.

## Data availability

Datasets related to this article can be found at http://dx.doi.org/10.17632/f8frrskzpz.1, an open-source online data repository hosted at Mendeley Data (Lovelace et al., 2021).

## Acknowledgements

We would like to thank the many volunteers, students, conservators and researchers who contributed to the excavation, preparation, curation, and study of NTC over the last four and a half decades. Our sincere appreciation to our colleagues xx for supportive dialog and guidance. We thank Jason Ballensky, Edmund Tucker, Ian Chechet, Ken Stahley, and Hans Bodenhamer for their efforts in surveying NTC. Thanks to Ryan Gardner for assistance in measuring sections and thoughtful discussion of observations. We would also like to thank Megan Sims (University of Kansas’ Vertebrate Paleontology Collec) for her help accessing information in the NTC database, and Emma Hendersen for helping collate archived data at the Oxford Radiocarbon Accelerator Unit. This project was made possible by the Wyoming Bureau of Land Management permit number PA-13-WY-207 to JAM, and we would like to thank B. Breithaupt and G. Hurley of the BLM for their assistance. This project was funded by Des Moines University (JAM), National Geographic Society Grant NGS 9479-14 to JAM, NSF EAR/SGP 1425059 to JAM, NSF EAR/SGP 1945013 to JLM, a Cave Conservancy grant to JAM and CR. Any use of trade, firm, or product names is for descriptive purposes only and does not imply endorsement by the U.S. Government.

## Supplementary Information: Appendix

**Figure 1.**
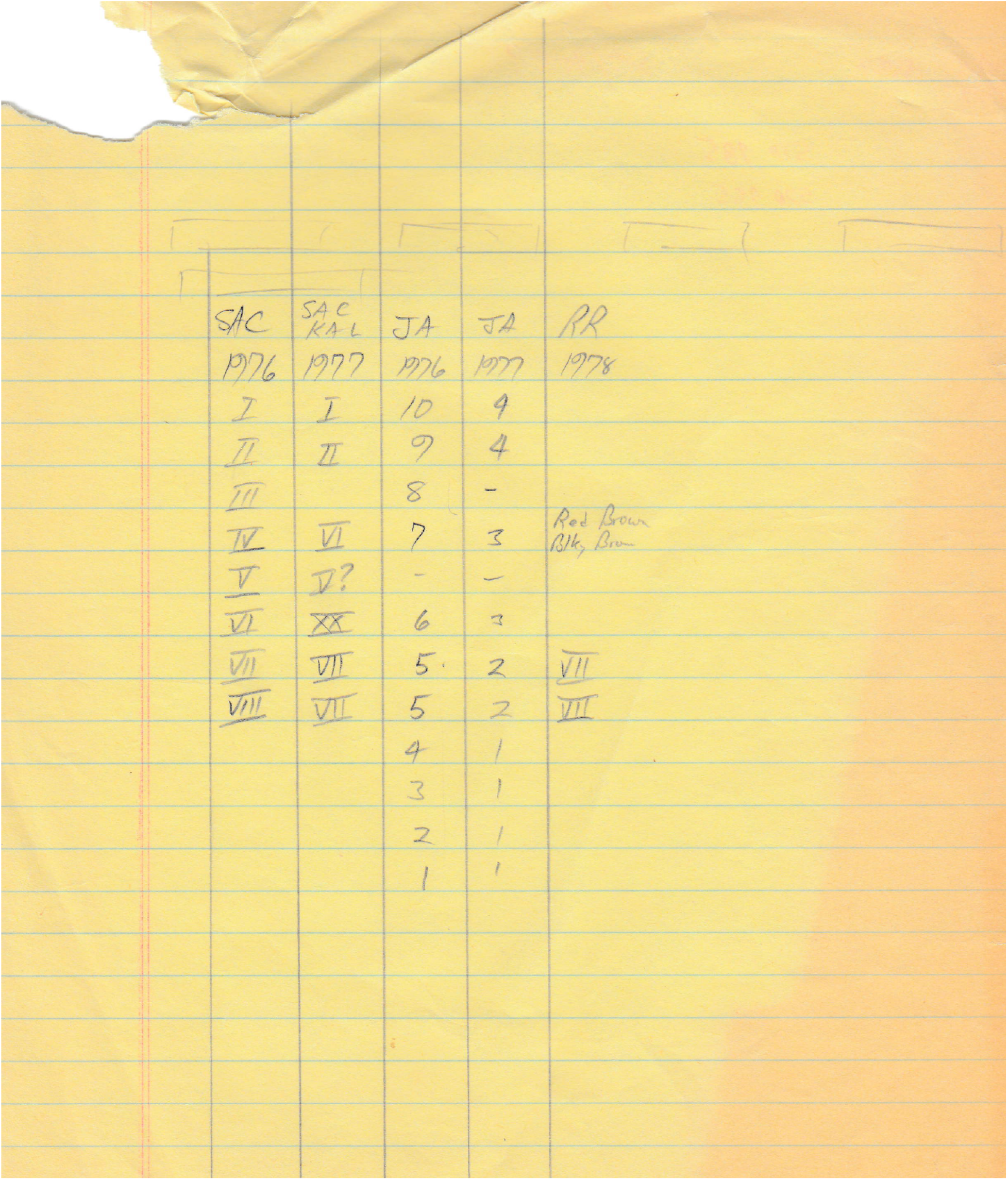
Hand-written correlations of early ‘zone’ and modified ‘zone’ nomenclature with that of Albanese’s (1976) nomenclature, Albanese’s 1977 written modifications, and a final partial system (RR) was not included in this study. SAC = Stephen A. Chomko; KAL = Kerry A. Lippencott; JA = John Albanese; RR = Randi Reppen.

**Table S1.** A List of specimens collected in the Lower Chamber in 2014 and 2016.

| Collection Year | UW_Specimen # | Taxon | ID element |
| --- | --- | --- | --- |
| 2014 | 50603 | Aves | coracoid |
| 2014 | 50604 | Aves | coracoid |
| 2014 | 50605 | Aves | coracoid |
| 2014 | 50606 | Aves | humerus |
| 2014 | 50607 | Aves | humerus |
| 2014 | 50608 | Aves | humerus |
| 2014 | 50609 | Aves | humerus |
| 2014 | 50610 | Aves | ulna |
| 2014 | 50611 | Aves | ulna |
| 2014 | 50612 | Aves | ulna |
| 2014 | 50613 | Aves | femur |
| 2014 | 50614 | Aves | femur |
| 2014 | 50615 | Aves | femur |
| 2014 | 50616 | Aves | femur |
| 2014 | 50617 | Aves | femur |
| 2014 | 50618 | Aves | femur |
| 2014 | 50619 | Aves | tibiotarsus |
| 2014 | 50620 | Aves | tibiotarsus |
| 2014 | 50628 | cf. Bison sp. | atlas |
| 2014 | 50629 | Bison sp. | lunar |
| 2014 | 50630 | Bison sp. | proximal phalanx |
| 2014 | 50631 | Bison sp. | middle phalanx |
| 2014 | 50632 | Bison sp. | distal phalanx |
| 2014 | 50633 | Bison sp. | distal phalanx |
| 2014 | 50694 | ?Bison sp. | cervical vertebra |
| 2014 | 50695 | ?Bison sp. | sacral vertebra |
| 2014 | 50696 | ?Bison sp. | thoracic vertebra |
| 2014 | 50635 | Ovis canadensis | metatarsal |
| 2014 | 50636 | Ovis canadensis | atlas |
| 2014 | 50637 | Ovis canadensis | ulna |
| 2014 | 50638 | Ovis canadensis | astragalus |
| 2014 | 50639 | Ovis canadensis | lunar |
| 2014 | 50640 | ?Bovidae | dentary |
| 2014 | 50641 | ?Bovidae | radius |
| 2014 | 50644 | cf. Canis lupus | proximal phalanx |
| 2014 | 50697 | cf. Canis lupus | cervical vertebra |
| 2014 | 50645 | Vulpes sp. | tibia |
| 2014 | 50648 | Mustela sp. | dentary |
| 2014 | 50650 | Lepus | humerus |
| 2014 | 50652 | Sylvilagus | humerus |
| 2014 | 50653 | Sylvilagus | radius |
| 2014 | 50660 | Lagomorpha | innominate |
| 2014 | 50676 | Equus sp. | femur |
| 2014 | 50698 | cf. Equus sp. | thoracic vertebra |
| 2014 | 50681 | Neotoma sp. | cranium |
| 2014 | 50699 | Mammalia | cervical vertebra |
| 2014 | 50700 | Mammalia | caudal vertebra |
| 2014 | 50701 | Mammalia | rib |
| 2014 | 50702 | Mammalia | rib |
| 2014 | 50703 | Mammalia | rib |
| 2016 | 53920 | Bison sp. | tooth |
| 2016 | 53949 | Canis sp. | metatarsal IV |

### 2.0 ^14^C analytical methods

#### OxA Determinations 2001-2005 Methods

AMS radiocarbon preparation procedures generally followed Brock et al. (2010). Bone or tooth fragments were sampled by using a tungsten carbide drill to generate coarse powder, which then underwent an acid-base-acid pretreatment of 0.5 M hydrochloric acid, followed by 0.1 M sodium hydroxide, and then another pretreatment of 0.5 M hydrochloric acid - with thorough rinsing using ultrapure water between each reagent. Collagen was then gelatinized at 75 C for 20 hr. Residual solids were filtered out prior to being ultrafiltered to remove the >30-kDa fraction. This fraction was freeze-dried and graphitized for AMS analysis.

#### LLNL Determinations 2015-2018 Methods

AMS radiocarbon preparation procedures generally followed Brown et al. (1988) and Bronk Ramsey et al. (2004). Decalcification with 0.25 N HCl was used to obtain collage that was then gelatinized at 58 °C for 16 h. Residual solids were filtered out prior to being ultrafiltered to remove the >30-kDa fraction. This fraction was freeze-dried and graphitized for AMS analysis. Dates were calibrated using OxCal v.4.4.3 (Bronk Ramsey, 2021), using the IntCal20 calibration curve (Reimer et al., 2020).

#### KCCAMS Determinations 2015-2018 Methods

AMS radiocarbon sample preparation largely follows Brown et al. (1988) and Bronk Ramsey et al. (2004). Samples were dissolved in 0.5 N HCl over 2 weeks (acid changed 3 times); bone chunks remained after dissolution. Filter pre-cleaning follows (Bronk Ramsey, 2004). Residual acid was removed and collagen was rinsed (Mili-Q water) prior to adding 10 mL of NaOH (removing humic acid). The NaOH was removed and collagen was rinsed (Mili-Q water) prior to adding 0.1 N HCl. Samples were gelatinized at 58 °C for 16 h. Unwound collagen was pre-filtered to remove residual solids, and then ultrafiltered to obtain >30-kDa fraction. The retantate was then freeze-dried and graphitized for AMS analysis. Dates were calibrated using OxCal v.4.4.3 (Bronk Ramsey, 2009), using the IntCal20 calibration curve (Reimer et al., 2020).

